# Gut-innervating nociceptor neurons protect against enteric infection by modulating the microbiota and Peyer’s patch microfold cells

**DOI:** 10.1101/580555

**Authors:** Nicole Y. Lai, Melissa A. Musser, Felipe A. Pinho-Ribeiro, Pankaj Baral, Pingchuan Ma, David E. Potts, Zuojia Chen, Donggi Paik, Salima Soualhi, Hailian Shi, Aditya Misra, Kaitlin Goldstein, Kisha N. Sivanathan, Amanda Jacobson, Antonia Wallrapp, Valentina Lagomarsino, Vijay K. Kuchroo, Roni Nowarski, Michael N. Starnbach, Neeraj K. Surana, Dingding An, Chuan Wu, Jun R. Huh, Meenakshi Rao, Isaac M. Chiu

## Abstract

Gut-innervating nociceptor sensory neurons respond to noxious/tissue-damaging stimuli by initiating protective responses and releasing mediators that regulate tissue inflammation, gastrointestinal secretion, and motility. The role of nociceptors in host defense against enteric pathogens is unclear. Here, we found that gut-extrinsic nociceptor neurons are critical in protecting the host against *Salmonella typhimurium* (STm) infection. Nociceptors responded to STm by releasing the neuropeptide calcitonin gene-related peptide (CGRP). Targeted depletion of Nav1.8 and TRPV1 neurons from gut-extrinsic dorsal root ganglia and vagal ganglia increased STm colonization, invasion, and dissemination. Nociceptors regulated the gut microbiota at homeostasis, specifically segmented filamentous bacteria (SFB) levels in the ileum, which protected against STm by colonization resistance. Nociceptors also regulated the density of microfold epithelial cells in the Peyer’s patch via CGRP to limit points of entry for STm invasion into host tissues. Understanding how host sensory neurons crosstalk with pathogenic bacteria may impact treatments for enteric infections.

**HIGHLIGHTS:**

- Nav1.8 and TRPV1 nociceptors defend against *Salmonella typhimurium* (STm) infection
- Nociceptors shape the gut microbiota and SFB levels which resist pathogen colonization
- Nociceptors suppress Peyer’s patch microfold cell density to limit pathogen invasion
- Neurons sense STm and release CGRP to modulate microfold cells and host defense

## INTRODUCTION

The gastrointestinal (GI) tract is one of the most heavily innervated organs in the body (Furness et al., 2009). Gut-innervating sensory neurons detect diverse stimuli, including mechanical stretch, dietary products, chemical and inflammatory stimuli (Blackshaw et al., 2007; Boesmans et al., 2011; Chavan et al., 2017; Holzer, 2002). Activation of these sensory neurons modulates gut physiological processes including satiety, pain, digestion, peristalsis, and fluid secretion in order to maintain gut homeostasis (Margolis et al., 2016; Van Der Zanden et al., 2009; Veiga-Fernandes and Pachnis, 2017; Yoo and Mazmanian, 2017).

Nociceptor neurons are specialized sensory neurons that detect harmful stimuli including heat, noxious chemicals, and inflammatory mediators (Basbaum et al., 2009; Benarroch, 2015; Woolf and Ma, 2007). Gut-innervating nociceptors initiate neural reflexes and sensations that protect the host from further damage, including visceral pain, nausea, and vomiting (Blackshaw and Gebhart, 2002; Di Giovangiulio et al., 2015; Grundy et al., 2018). However, whether gut-innervating nociceptor neurons crosstalk with microbes and immune cells to regulate mucosal host defenses is not well understood.

One of the major threats to gut homeostasis is invasion by enteric pathogens. In this study, we examined the role of nociceptors in enteric host defense against the Gram-negative bacterial pathogen *Salmonella enterica* serovar Typhimurium (STm). Oral infections of STm causes typhoid-like disease in mice and gastroenteritis in humans (Coburn et al., 2007). Despite antibiotic treatments, *Salmonella* infections still represent a massive clinical burden, causing 94 million cases and 155 thousand deaths worldwide each year (Majowicz et al., 2010). Given the rise of antibiotic-resistant *Salmonella* strains (Andrews et al., 2018), there is a critical need to enhance our understanding of host-pathogen interactions to explore better host-directed treatments for enteric infections (Behnsen et al., 2015; Broz et al., 2012; Santos et al., 2009). During oral infections of mice, STm travels through the GI tract to transiently colonize the distal small intestine. After entering the ileum, STm invades host tissue by crossing into Peyer’s patches, then spreads via efferent lymphatics to the mesenteric lymph nodes. Subsequently, STm disseminates and replicates in peripheral organs, including the spleen and liver, to cause systemic disease (Monack et al., 2000; Vazquez-Torres et al., 1999).

An important host protection mechanism that limits intestinal colonization of pathogens, including STm, is resistance mediated by commensal microbes of the GI tract (Kamada et al., 2013; Stecher and Hardt, 2011). Resident commensals compete for nutrients and secrete metabolites/toxins that can directly counteract incoming pathogens (Baumler and Sperandio, 2016; Buffie and Pamer, 2013), as well as indirectly stimulate host anti-bacterial mucosal responses that impact pathogen survival in the GI tract (Abraham and Medzhitov, 2011; Blander et al., 2017; Littman and Pamer, 2011). STm utilizes several virulence factors encoded on *Salmonella* Pathogenicity Islands (SPI-1 and SPI-2) to robustly colonize the GI tract and establish a niche within the host (McGhie et al., 2009; Thiennimitr et al., 2012).

Following colonization of the distal small intestine, the second step of STm infection is invasion of host tissues. Although different entry routes by STm into host tissues have been described, such as invasion through enterocytes and luminal uptake via transepithelial processes from phagocytic cells (Kulkarni et al., 2018; Rescigno et al., 2001; Sansonetti, 2004), STm’s predominant route of entry into host tissues is through invasion of microfold (M) cells in the Peyer’s patches (Jang et al., 2004; Jepson and Clark, 2001; Jones et al., 1994). M cells are specialized antigen-sampling epithelial cells overlaying the follicles in Peyer’s patch and they transcytose antigens from the lumen to underlying phagocytes to initiate mucosal immunity (Mabbott et al., 2013; Ohno, 2016; Williams and Owen, 2015). Enteric microbial pathogens, including STm, *Vibrio cholerae*, poliovirus, and prions, utilize M cells as key entry points to invade and disseminate from the gut to cause disease (Donaldson et al., 2016a; Miller et al., 2007; Sansonetti and Phalipon, 1999). STm has also been found to transdifferentiate follicle-associated epithelial cells into M cells to facilitate its invasion of the small intestine (Savidge et al., 1991; Tahoun et al., 2012).

In this study, we demonstrate that nociceptors play a major role in protecting the host against STm infection. Through molecular targeting of Nav1.8+ and TRPV1+ nociceptors, we found that depletion of these neurons exacerbated disease by increasing STm colonization of the ileum, invasion into Peyer’s patches, and dissemination to extraintestinal organs. We identified TRPV1+ DRG nociceptors as the critical neural subset mediating host protection and showed that DRG neurons responded directly to STm by releasing the neuropeptide CGRP. Nociceptors modulated the composition of the small intestine microbiota to promote host protection against STm colonization, specifically through the ileal resident microbe Segmentous Filamentous Bacteria (SFB). Nociceptors also regulated the density of M cells in ileal Peyer’s patches to protect against STm invasion and dissemination. These findings demonstrate that the gut-innervating nociceptors play an active role in maintaining gut homeostasis and host protection against pathogens. Future therapies that target nociceptor neurons could lead to enhanced host mucosal protection and treatments for enteric pathogen infections.

## RESULTS

### Nociceptor neurons mediate host protection against *Salmonella* infection

The sensory innervation of the GI tract includes gut-extrinsic neurons whose cell bodies reside within the dorsal root ganglia (DRG) and nodose/jugular vagal ganglia (VG), and intrinsic primary afferent neurons whose cell bodies reside within the myenteric and submucosal plexus of the gut (Fig. 1A) (Furness et al., 2009; Uesaka et al., 2016). DRG and vagal nociceptor neurons mediate the sensations of visceral pain and nausea/vomiting, respectively (Blackshaw and Gebhart, 2002). However, whether nociceptors protect the GI tract against enteric pathogens is unclear. To study the role of nociceptors in enteric host defense, we infected mice orally with SL1344, a wildtype STm strain that colonizes the distal small intestine, invades the Peyer’s patches, and spreads to peripheral organs including the liver and spleen (Monack et al., 2000).

**Figure 1.**
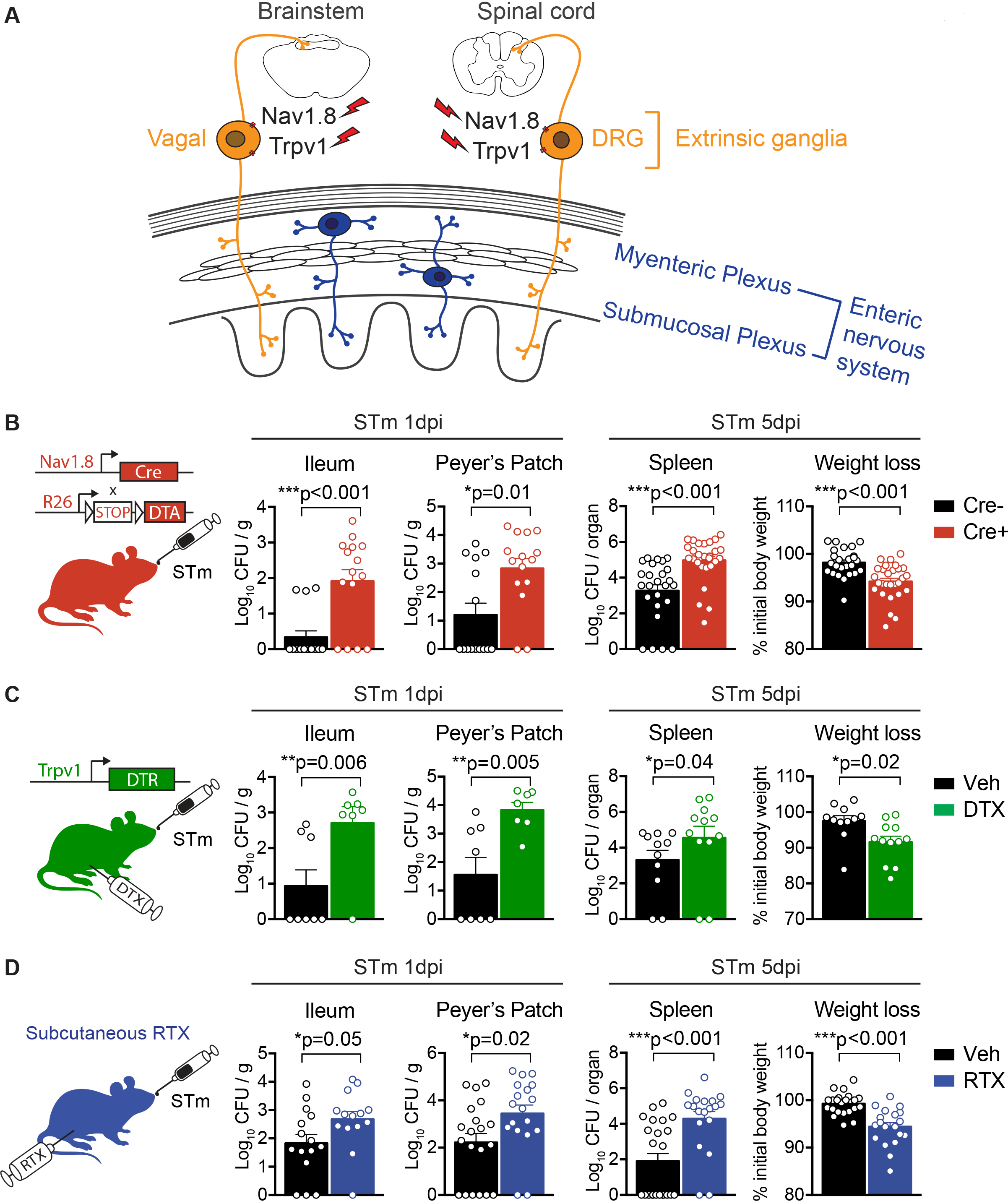
Nociceptor neurons mediate host protection against *Salmonella* infection. **(A)** Schematic diagram showing sensory neuron innervation of the gastrointestinal tract. Extrinsic sensory ganglia include the nodose/jugular vagal ganglia and dorsal root ganglia (DRG), which house sensory neurons that transduce signals from different layers of the gut and project to the brainstem and spinal cord, respectively. We used genetic and pharmacological strategies to specifically target Nav1.8 and TRPV1 nociceptor neurons within extrinsic ganglia. Within the gut, the myenteric plexus and submucosal plexus house intrinsic primary afferent neurons that comprise the sensory arm of the enteric nervous system. **(B)** *Nav1.8-Cre*^+^/*DTA* (Cre+) mice genetically ablated of Nav1.8-lineage neurons and *Nav1.8-Cre*^−^/*DTA* (Cre-) control littermates were orally infected with *Salmonella typhimurium* (STm) and sacrificed at 1 or 5 days post-infection (dpi) to examine STm burden in different tissues, and weight loss. At 1 dpi, data are pooled from 3 experiments with Cre- (n=15) and Cre+ (n=15) mice infected with 0.8-1.1 × 10^7^ colony forming units (CFU) STm. At 5 dpi, data are pooled from 5 experiments with Cre- (n=25) and Cre+ (n=28) mice infected with 1.1-1.6 × 10^7^ CFU STm. **(C)***Trpv1-DTR* mice injected with diphtheria toxin (DTX) or vehicle (PBS) were orally infected with STm and sacrificed at 1 or 5 dpi to examine STm burden in different tissues and weight loss. At 1 dpi, data are pooled from 2 experiments from vehicle (n=8) and DTX (n=7) treated mice infected with 0.8-1.8 × 10^7^ CFU STm. At 5 dpi, data are pooled are from 2 experiments from vehicle (n=11) and DTX (n=12) treated mice infected with 0.9-1.2 × 10^7^ CFU STm. **(D)**Wildtype mice injected subcutaneously with resiniferatoxin (RTX; chemically ablated of TRPV1 neurons) or vehicle (Veh) were orally infected with STm and sacrificed at 1 or 5 dpi to examine STm burden in different tissues and weight loss. At 1 dpi, data are pooled from 4 experiments from vehicle (n=15-20) and RTX (n=14-18) treated mice infected with 0.7-1.8 × 10^7^ CFU STm. At 5 dpi, data are pooled from 4 experiments from vehicle (n=23) and RTX (n=19) treated mice infected with 1.2-1.8 × 10^7^ CFU STm. Statistical analysis: **(B-D)** Each circle represents an individual animal. Mann-Whitney tests for CFU analysis. Unpaired t-tests for weight loss analysis. *p<0.05, **p<0.01, ***p<0.001. Error bars are mean ± SEM. See also Figure S1.

We first established molecular strategies to selectively target nociceptor neurons. Single cell RNA-sequencing data was recently published for peripheral and central nervous system cell-types, including enteric neurons, DRG neurons, and glia (Zeisel et al., 2018). Mining this data, we found that *Scn10a* (encoding the nociceptor-enriched sodium channel Nav1.8) and *Trpv1* (encoding the TRPV1 ion channel that detects noxious heat) genes are highly expressed by DRG neurons, but are absent in enteric neurons of the gut (**Fig. S1A**). We reasoned that experimental strategies targeting Nav1.8+ or TRPV1+ neurons would target gut-extrinsic nociceptor neurons without affecting intrinsic enteric neurons of the gut (Fig. 1A).

Nav1.8 is a tetrodotoxin-resistant voltage-gated sodium channel selectively expressed by nociceptors that can be sensitized by inflammatory mediators including cytokines and prostaglandins (Lai et al., 2004). We targeted Nav1.8+ nociceptors by breeding *Nav1.8-Cre*^+/−^ mice with *Rosa-Diphtheria toxin A* (*DTA*) reporter mice, where Cre-mediated DTA expression leads to developmental ablation of Nav1.8-lineage nociceptors in *Nav1.8-Cre*^+^/*DTA* mice (hereafter called *Nav1.8-Cre/DTA* or Cre+) compared to *Nav1.8-Cre*^−^/*DTA* control littermates (Cre-) (Abrahamsen et al., 2008). Immunostaining of DRG and vagal ganglia from *Nav1.8-Cre/DTA* mice showed reduced proportions of both TRPV1+ and CGRP+ nociceptor neurons compared to control littermates (**Fig. S1C-D**). Conversely, the proportion of NF200+ neurons, representing large diameter A-fibers, was increased in DRG and vagal ganglia from *Nav1.8-Cre/DTA* mice (**Fig. S1C-D**). These neuron proportion changes reflect a previous study showing that Nav1.8-lineage neurons mediate cold, mechanical and inflammatory pain in mice (Abrahamsen et al., 2008).

Following oral infection with STm, *Nav1.8-Cre/DTA* mice exhibited significantly increased STm levels in the small intestine ileum (p<0.001) and increased STm invasion into Peyer’s patches (p=0.01) compared to control littermates at 1 day post-infection (dpi) (Fig. 1B). At this same time-point, STm levels were also increased in the cecum and mesenteric lymph nodes (LNs) of *Nav1.8-Cre/DTA* mice compared to control littermates (**Fig. S1B**). We next examined STm dissemination to systemic sites at a later time point of 5 dpi. There was significantly greater STm burden in the spleen (p<0.001) and liver (p<0.001) of *Nav1.8-Cre/DTA* mice compared to control littermates (Fig. 1B and **Fig. S1B**). *Nav1.8-Cre/DTA* mice also exhibited greater body weight loss (p<0.001) compared to control littermates (Fig. 1B). Therefore, Nav1.8+ neurons control early invasion and later systemic dissemination of STm in mice.

TRPV1 (transient receptor potential vanilloid 1) is a nociceptive ion channel that responds to heat (>45°C), protons, and capsaicin (Julius, 2013). TRPV1+ neurons are required for the detection of noxious thermal sensation (Mishra et al., 2011) and inflammatory pain (Mishra and Hoon, 2010). TRPV1 and Nav1.8 largely overlap in gene expression, but not completely, in the DRG (Amaya et al., 2000). Recent data from tracing of colonic sensory neurons show that *Trpv1* and *Scn10a* (Nav1.8) gut-innervating neural subsets mostly overlap, but did also include some distinct populations (https://hockley.shinyapps.io/ColonicRNAseq). Therefore, it was informative to determine each of their roles in host defense.

To determine whether TRPV1+ neurons are involved in enteric host defense, we utilized *Trpv1-DTR* mice, which express the human diphtheria toxin receptor (DTR) under the *Trpv1* promoter (Pogorzala et al., 2013). Injections of diphtheria toxin (DTX) into *Trpv1-DTR* mice have been shown to specifically ablate TRPV1+ expressing neurons in the DRG and vagal ganglia (Baral et al., 2018; Trankner et al., 2014). Following oral infection with STm, *Trpv1-DTR* mice treated with DTX exhibited significantly increased STm burden in the ileum (p=0.006), Peyer’s patches (p=0.005), and mesenteric LNs (p=0.01) at 1 dpi compared to vehicle-injected control mice (Fig. 1C and **Fig. S1E**). TRPV1+ neuron-ablated mice also showed significantly increased STm dissemination to the spleen (p=0.04) and liver (p=0.03), and increased weight loss (p=0.02) at 5 dpi (Fig. 1C and **Fig. S1E**). As an additional strategy, we chemically ablated TRPV1+ nociceptors by injecting mice with resiniferatoxin (RTX), a potent TRPV1 agonist that leads to pharmacological denervation of these neurons (Liu et al., 2010; Szolcsanyi et al., 1991). Following RTX treatment, mice were rested for 4 weeks before oral infections of STm. RTX-treated mice displayed significantly greater STm load in the ileum (p=0.05), Peyer’s patches (p=0.02), and mesenteric LNs (p=0.02) at 1 dpi (Fig. 1D and **Fig. S1F**). RTX-treated mice had greater STm dissemination to the spleen (p<0.001) and liver (p<0.001), as well as greater weight loss (p<0.001) at 5 dpi (Fig. 1D and **Fig. S1F**). We confirmed that RTX-treated mice had significantly decreased proportions of TRPV1+ and CGRP+ neurons in DRG and vagal ganglia compared to vehicle-treated mice (**Fig. S1G-H**).

Therefore, we found that nociceptor neurons play a critical role in protecting against STm infection. Using three different approaches, one that targeted Nav1.8+ nociceptors (*Nav1.8-Cre/DTA*) and two that targeted TRPV1+ nociceptors (*Trpv1-DTR*, RTX treatment), we found that all three strategies significantly worsened outcomes of infection, including increased STm colonization of the ileum, invasion of Peyer’s patches, and dissemination to peripheral organs.

### Gut-extrinsic TRPV1+ DRG neurons protect against *Salmonella* infection

Given that nociceptor neurons reside in both the DRG and vagal ganglia (Fig. 1A), we aimed to define the key neuronal subsets mediating enteric host defense against STm. We analyzed whether Nav1.8 or TRPV1-targeting strategies were specific to extrinsic sensory neurons, and whether gut-intrinsic enteric neurons were impacted in these mice. Using qRT-PCR of tissues from the DRG, vagal ganglia, and myenteric plexus/longitudinal muscle preparations of the ileum and colon, we observed complete loss of *Scn10a* transcripts in extrinsic ganglia of *Nav1.8-Cre/DTA* mice (Fig. 2A), and complete loss of *Trpv1* transcripts in extrinsic ganglia of RTX-treated mice (Fig. 2B). *Trpv1* and *Scn10a* transcripts were below the limit of detection in myenteric plexus preparations from these mice. Conversely, *Hand2*, a transcription factor involved in terminal differentiation of enteric neurons (D’Autreaux et al., 2007), was highly expressed in the myenteric plexus and did not differ between nociceptor-ablated and control mice **(Fig. S2A-B)**.

**Figure 2.**
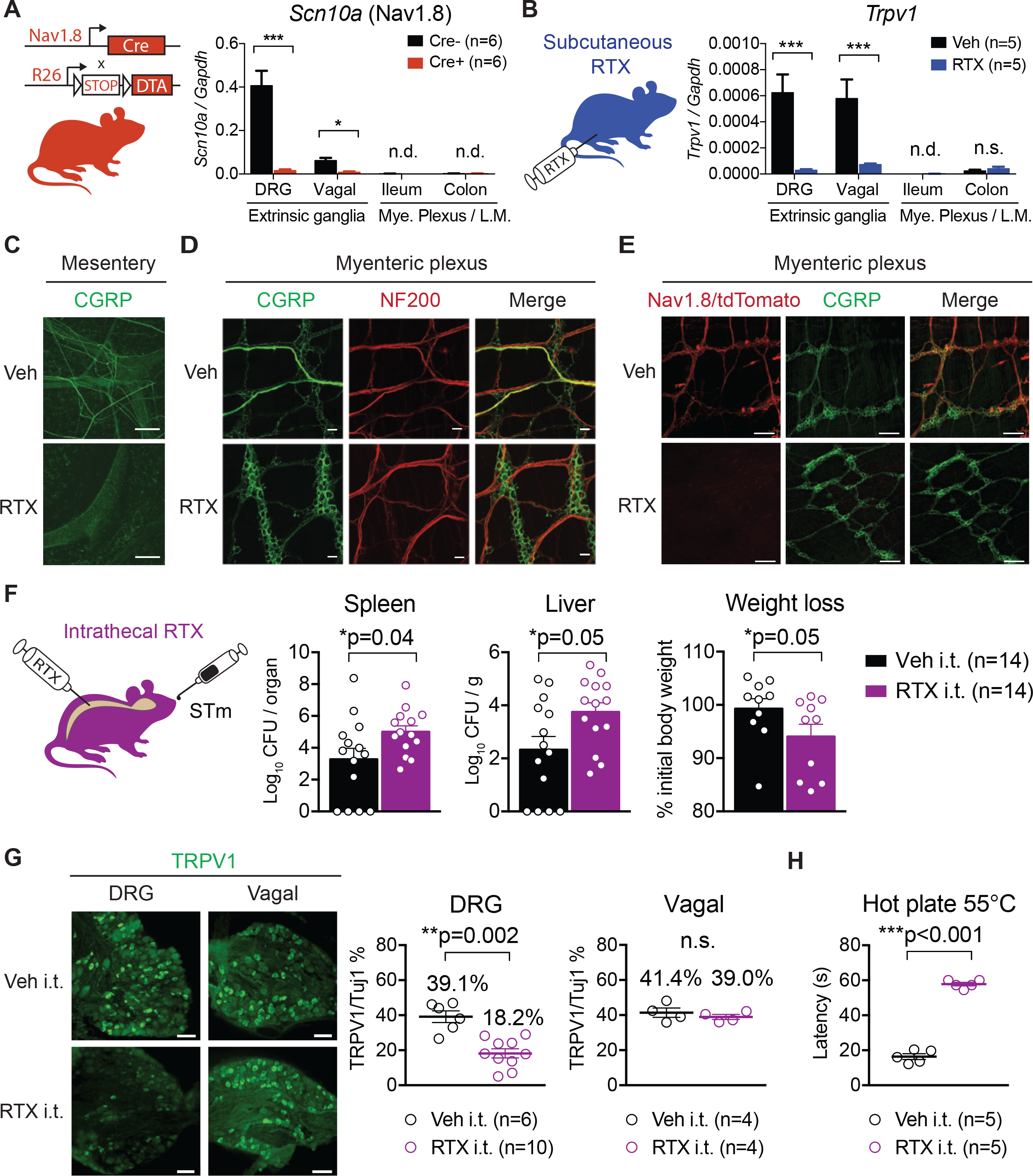
Spinal TRPV1+ nociceptor neurons are a major neural subset that protect against *Salmonella* infection. **(A)** qRT-PCR of *Scn10a* transcripts (encoding Nav1.8) from DRG, vagal ganglia, ileum, and colon myenteric plexus/longitudinal muscle tissues of *Nav1.8-Cre/DTA* mice and control littermates (n=6 mice/group). **(B)** qRT-PCR of *Trpv1* transcripts from DRG, vagal ganglia, ileum and colon tissues of vehicle-treated and RTX-treated mice (n=5 mice/group). **(C)** The mesentery was immunostained for the neuropeptide CGRP in vehicle and RTX-treated mice. The mesentery contains afferent and efferent nerve fibers to the gastrointestinal tract, Scale bar, 100 um. **(D)** Myenteric plexus was stained for CGRP and neurofilament 200 (marker for large diameter A-fibers). RTX-treated mice exhibited a loss of thick CGRP nerve tracts carrying extrinsic fibers to the myenteric plexus. Within the myenteric plexus, CGRP+ intrinsic enteric neurons remained intact. Scale bar, 25 um. **(E)** *Nav1.8-Cre/tdTomato* mice treated with RTX exhibited a loss of tdTomato+ extrinsic sensory fibers compared to vehicle-treated mice, while CGRP+ intrinsic enteric neurons remained intact in the myenteric plexus. Scale bar, 100 um. **(F)** Wildtype mice injected intrathecally with RTX or vehicle were orally infected with STm and sacrificed at 5 dpi to examine STm dissemination and weight loss. Data are pooled from 3 experiments with an infection dose of 1.2-1.8 × 10^7^ CFU from vehicle (n=14) and RTX (n=14) intrathecally injected mice. **(G)** Quantification of the proportions of TRPV1+ neurons out of total Tuj1+ neurons in lumbar DRGs and vagal ganglia of intrathecally injected mice. Data are from vehicle (n=3) and RTX (n=3) intrathecally injected mice. Scale bar, 50 um. **(H)** Hot plate latency for vehicle and RTX intrathecally injected mice (n=5 mice/group). Statistical analysis: **(A-B)** Two–way ANOVA and Sidak post-tests for qRT-PCR. **(F)** Mann-Whitney tests for CFU analysis. Unpaired t-test for weight loss analysis. **(G, H)** Unpaired t-tests for DRG and vagal ganglia immunostaining analysis, and hot plate analysis. *p<0.05, **p<0.01, ***p<0.001. Error bars are mean ± SEM. See also Figure S2.

We next examined extrinsic sensory nerve fibers entering the GI tract. Staining of the mesentery demonstrated an absence of extrinsic nerve fibers expressing CGRP, a neuropeptide found in peptidergic nociceptors, in RTX-treated mice compared to vehicle-treated mice (Fig. 2C). We also observed a loss of thick CGRP+ nerve tracts carrying extrinsic fibers to the myenteric plexus in RTX-treated mice, though intrinsic CGRP+ enteric neurons were still present (Fig. 2D). RTX treatment also diminished fluorescently labeled Nav1.8+ nerve fibers in the myenteric plexus of *Nav1.8-Cre*/*tdTomato* reporter mice (Fig. 2E). We next performed immunostaining analysis of intrinsic enteric neuron populations in the myenteric plexus. The total number of neurons expressing HuD+ (a pan-neuronal marker) did not differ in *Nav1.8-Cre/DTA* or RTX-treated mice compared to their controls **(Fig. S2C-H)**. The proportion of calretinin+ cells, which marks a subset of enteric neurons, and neuronal nitric oxide synthase (nNOS+), which marks nitrergic neurons, were also unchanged in the myenteric plexi of nociceptor-ablated mice compared to their controls **(Fig. S2C-H)**. These data showed that our targeting strategies led to selective loss of gut-extrinsic sensory neurons, while leaving gut-intrinsic neurons and the overall architecture of the network of myenteric neurons intact. Of note, such molecular strategies that selectively target extrinsic sensory neurons could be useful for the GI field beyond our study.

We next focused on determining whether gut-extrinsic DRG nociceptors specifically contributed to enteric host defense. Intrathecal injections of RTX into the spinal cord led to depletion of TRPV1+ neurons in the DRG, while leaving TRPV1+ neurons in the vagal ganglia unaffected (Fig. 2G). RTX-intrathecally injected mice had significantly delayed nocifensive responses on a hot plate assay, which reflected the loss of noxious heat sensation mediated by spinal TRPV1+ neurons (Fig. 2H). We next infected RTX- and vehicle-intrathecally injected mice with STm. Specific ablation of TRPV1+ DRG neurons led to significantly higher STm dissemination to the spleen (p=0.04) and liver (p=0.05), and increased weight loss (p=0.05) at 5 dpi (Fig. 2F), which phenocopied earlier data from RTX subcutaneous injections. Intrathecal injections of RTX are unlikely to affect TRPV1+ cells outside the central nervous system, thus ruling out effects on non-neuronal or immune cells in the periphery. These data demonstrated that gut-extrinsic TRPV1+ DRG neurons are a major neuronal subset that mediate enteric host defense against STm infection.

We sought to determine which step of STm infection was regulated by nociceptor neurons. Intraperitoneal infections of STm were performed to bypass invasion of the gut barrier to examine whether nociceptors modulate STm replication in its systemic phase. We did not detect differences in STm burdens in spleens and livers between vehicle and RTX-treated mice following intraperitoneal infection **(Fig. S2I)**. These data indicate that nociceptors do not regulate STm survival or replication in peripheral organs, and likely mediate host defense at earlier stages of infection prior to STm extraintestinal dissemination such as ileum colonization (Fig. 1B-D).

### Nociceptors regulate the small intestine ileum microbiota and maintain segmented filamentous bacteria (SFB)

The gut microbiota plays an important role in mediating colonization resistance to enteric bacterial infections, including STm (Baumler and Sperandio, 2016; Stecher et al., 2005), and in shaping GI mucosal immune responses (Blander et al., 2017; Littman and Pamer, 2011). The role of sensory neurons, specifically nociceptors, in shaping the gut microbiota has not been explored. Given that nociceptor-ablated mice had higher levels of STm in the ileum (Fig. 1B-D), we hypothesized that nociceptors may regulate microbial homeostasis, which could mediate colonization resistance against STm.

We performed analysis of gut bacterial communities using 16S rRNA gene sequencing in *Nav1.8-Cre/DTA* mice to determine whether nociceptors modulate the gut microbiota at homeostasis. We sampled the luminal contents and mucosal scrapings from different segments of the GI tract, including the duodenum, ileum, and colon of singly-housed *Nav1.8-Cre/DTA* mice (n=6) and control littermates (n=6) at 10 weeks of age (Fig. 3). We also examined feces from these mice at 4 weeks and 10 weeks of age. We noted a marked increase in bacterial diversity as measured by the Chao1 index and rarefaction curves in *Nav1.8-Cre/DTA* ileum lumen and mucosa samples compared to control littermate samples (Fig. 3A-B). By contrast, bacterial richness was broadly similar between *Nav1.8-Cre/DTA* and control littermates across the duodenum, colon, and feces. Principal component analysis (PCA) showed separation between *Nav1.8-Cre/DTA* and control mucosa and lumen samples from the ileum, whereas PCA of duodenum and colon samples from these same mice showed greater overlap (Fig. 3C). The major differences in the bacterial composition of the ileum mucosa from *Nav1.8-Cre/DTA* mice corresponded to decreased abundance of *Firmicutes* and increased abundance of *Bacteroidetes* at the phylum level compared to control mice (Fig. 3D). Within the *Firmicutes*, the most striking change was a reduction of *Clostridiaceae* family members in the ileum lumen and mucosa of *Nav1.8-Cre/DTA* mice (Red bar, Fig. 3E). This was largely accounted for at the genus level by significantly lower levels of segmented filamentous bacteria (SFB) (Bolotin et al., 2014; Ivanov et al., 2009) in *Nav1.8-Cre/DTA* mice (Fig. 4A-B).

**Figure 3.**
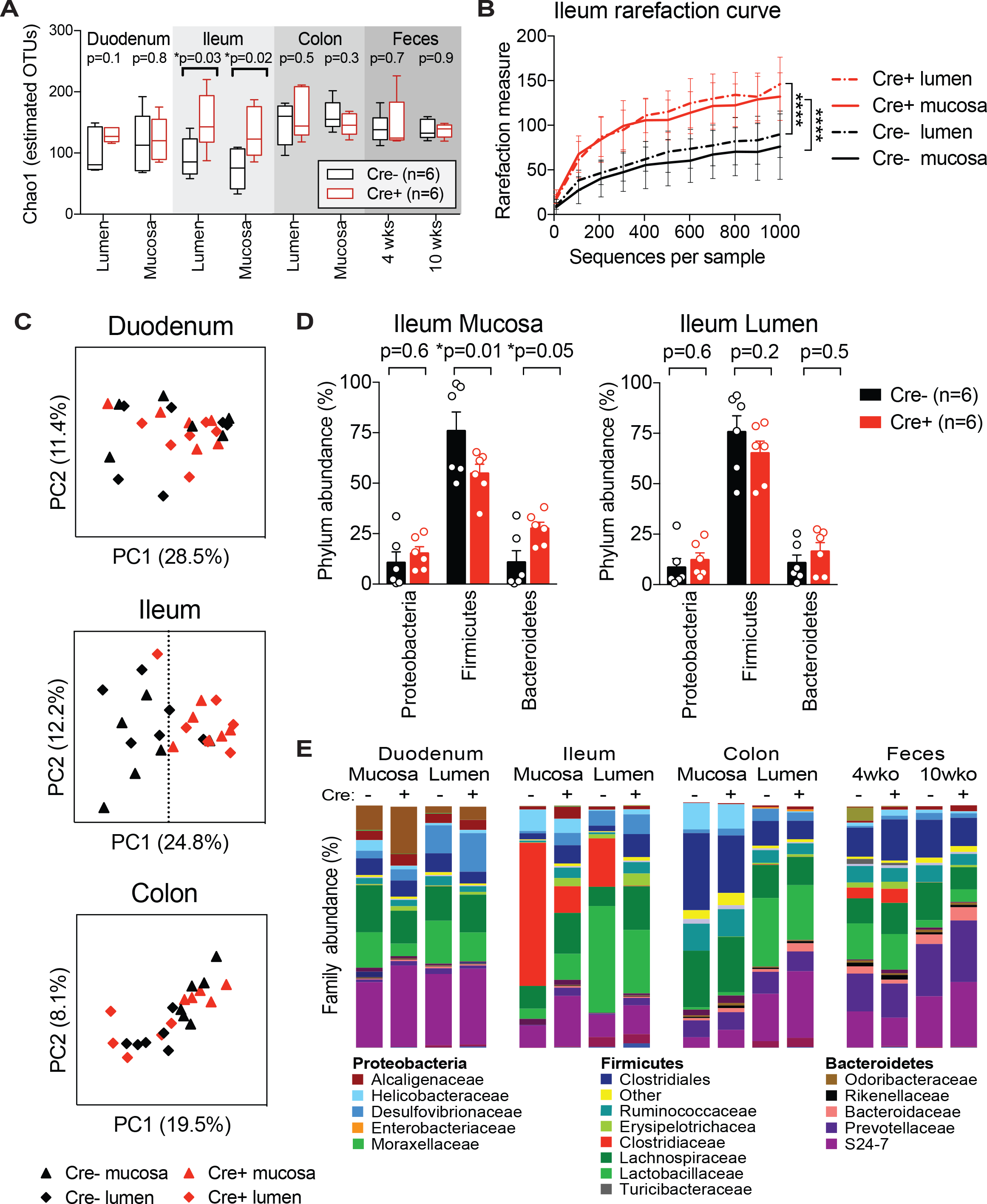
Nociceptor neurons regulate bacterial composition in the ileum. *Nav1.8-Cre/DTA* mice and control littermates (n=6 Cre- and n=6 Cre+ from 2 litters) were single-housed from weaning and feces were collected at 4 and 10 weeks of age. At 10 weeks of age, mice were sacrificed, then lumenal contents and mucosal scrapings from the duodenum, ileum, and colon were subjected to 16S rRNA sequencing. **(A)** Estimated bacterial diversity, as determined by Chao1 index, in *Nav1.8-Cre/DTA* mice and control littermates (n=6 mice/group). **(B)** Rarefaction curve using Chao1 index of ileum lumen and mucosa samples from *Nav1.8-Cre/DTA* mice and control littermates (n=6 mice/group). **(C)** Principal component analysis of duodenum, ileum, and colon samples from *Nav1.8-Cre/DTA* mice and control littermates (n=6 mice/group). **(D)** Bar charts showing the relative phylum abundance of ileum mucosa and lumen samples from *Nav1.8-Cre/DTA* mice and control littermates (n=6 mice/group). **(E)** Bar charts showing the relative family abundance of tissues from *Nav1.8-Cre/DTA* mice and control littermates (n=6 mice/group). Statistical analysis: **(A)** Multiple t-tests for Chao1 index. **(B)** One-way ANOVA repeated measures and Sidak post-tests for rarefaction curve. **(D)** Two-way ANOVA and Fisher’s post-test for phylum analysis. *p<0.05, ****p<0.0001. Error bars are mean ± SEM. See also Figure S3 and S4.

**Figure 4.**
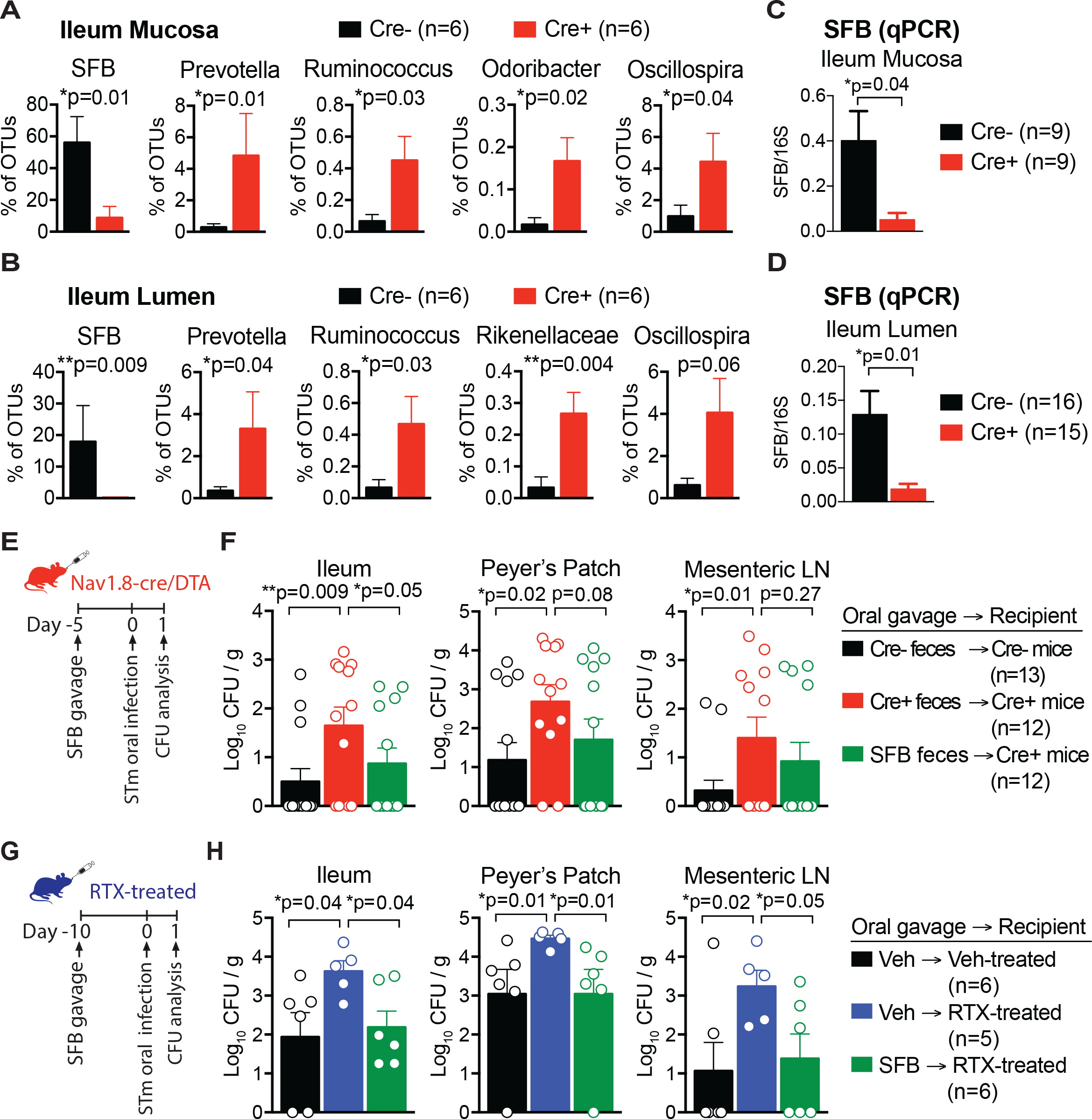
Nociceptor regulation of segmented filamentous bacteria in the ileum provides colonization resistance against *Salmonella* infections. **(A-B)** Differentially abundant microbes in the ileum mucosa (A) and ileum lumen (B) of *Nav1.8-Cre/DTA* mice and control littermates (n=6 mice/group). **(C and D)** qPCR of SFB levels in ileum mucosa (C) and ileum lumen (D) from *Nav1.8-Cre/DTA* mice and control littermates. Data are pooled across 7 litters for ileum mucosa (n=9 Cre- and n=9 Cre+) and ileum lumen (n=16 Cre- and n=15 Cre+) samples. **(E-F)** *Nav1.8-Cre/DTA* mice and control littermates were gavaged with feces from SFB-monocolonized germ-free mice, or their own feces as control. Five days after fecal transplantation, mice were orally infected with 0.76-1.08 × 10^7^ CFU STm and sacrificed at 1 dpi to analyze STm tissue burden. Data are pooled from Cre-mice gavaged with self-feces (n=12), Cre+ mice gavaged with self-feces (n=12), and Cre+ mice gavaged with SFB feces (n=12). **(G-H)** Vehicle-treated and RTX-treated mice were gavaged with feces from SFB-monocolonized germ-free mice, or with vehicle (water) as control. Ten days after SFB transplantation, mice were orally infected with 0.59 × 10^7^ CFU STm and sacrificed at 1 dpi to analyze STm tissue burden. Data are from vehicle-treated mice gavaged with vehicle (n=6), RTX-treated mice gavaged with vehicle (n=5), and RTX-treated mice gavaged with SFB feces (n=6). Statistical analysis: **(A-D)** Mann-Whitney tests for microbial differential abundance and SFB qPCR. **(F, H)** Non-parametric one-way ANOVA and Dunn’s post-tests for CFU analysis. *p<0.05, **p<0.01. Error bars are mean ± SEM. See also Figure S4.

In control mice, SFB made up a large proportion of ileum mucosal bacteria (56% of OTUs), whereas this level was significantly decreased in *Nav1.8-Cre/DTA* mice (9% of OTUs) (p=0.01) (Fig. 4A). Similarly, in the lumen of the ileum, control mice had higher levels of SFB (18% of OTUs) compared to *Nav1.8-Cre/DTA* mice (0.2% of OTUs) (p=0.009) (Fig. 4B). *Nav1.8-Cre/DTA* mice also displayed significantly increased levels of several microbial members (*Prevotella, Ruminococcus, Odoribacter, Rickenellaceae, Oscillospira*), which were minor components compared to SFB (Fig. 4A-B). Next, to confirm the 16S sequencing analysis, we analyzed SFB levels in *Nav1.8-Cre/DTA* mice using SFB-specific qPCR. Across 7 different litters of mice, we found significantly lower SFB levels in ileum lumen and mucosal samples from *Nav1.8-Cre/DTA* mice compared to control littermates (Fig. 4C-D).

These data indicate that nociceptors specifically control microbial homeostasis in the ileum relative to duodenum, colon, and feces by maintaining homeostatic levels of SFB. The small intestine is the major entry point of STm into host tissues (Jones et al., 1994), and within the small intestine, STm has been shown to preferentially invade the ileum rather than the jejunum (Rivera-Chavez et al., 2016). Thus, nociceptor maintenance of microbial homeostasis in the ileum could potentially regulate host defense against STm.

### Nociceptor regulation of SFB mediates resistance against STm infection

SFB has been shown to be protective against enteric bacterial pathogens including *Salmonella enteriditis, Citrobacter rodentium*, and *Escherichia coli* (Garland et al., 1982; Heczko et al., 2000; Ivanov et al., 2009). To test whether SFB could enhance host protection against STm infection in nociceptor-ablated mice, we gavaged *Nav1.8-Cre/DTA* mice with fecal pellets from SFB-monocolonized germ-free mice or their own feces as control. Mice were rested for five days to facilitate robust SFB colonization (Farkas et al., 2015), then were orally infected with STm and analyzed at 1 dpi (Fig. 4E). As observed previously, the ileums and Peyer’s patches of *Nav1.8-Cre/DTA* mice had higher STm levels compared to control littermates. SFB transplantation into *Nav1.8-Cre/DTA* mice significantly decreased STm levels in the ileum and Peyer’s patch at 1 dpi (Fig. 4F). We also performed an experiment examining STm burden at a later time point. At 5 dpi, STm dissemination to the liver and spleen was also reduced in SFB-gavaged mice **(Fig. S4A)**. Successful SFB colonization was confirmed by qPCR analysis of ileum mucosa from these mice **(Fig. S4B)**. These data indicated that SFB can restore STm host defense in Nav1.8+ neuron deficient mice.

We next asked whether SFB could confer similar host protection against STm in TRPV1 neuron-ablated mice. Vehicle-treated or RTX-treated mice were gavaged with vehicle or fecal pellets from SFB-monocolonized germ-free mice. Following SFB transplantation, mice were infected with STm and analyzed at 1 dpi (Fig. 4G). Transplantation of SFB into RTX-treated mice significantly reduced STm colonization of the ileum (p=0.04), invasion of Peyer’s patches (p=0.01), and dissemination to mesenteric LN (p=0.05), suggesting that SFB can restore host protection against STm in TRPV1+ neuron-ablated mice (Fig. 4H).

We next investigated if immune populations regulated by SFB may play a role in nociceptor-mediated host defense. Previous studies have shown that SFB attaches to the ileal mucosal lining and critically mediates the development of Th17 cells in mice (Atarashi et al., 2015; Gaboriau-Routhiau et al., 2009; Ivanov et al., 2009). We therefore measured Th17 cells (IL17A+CD4+TCRβ+) in the ileal lamina propria at homeostasis and after STm infection **(Fig. S4C)**. Our analysis did not detect differences in Th17 cell populations between *Nav1.8-Cre/DTA* and control littermates. We also did not detect differences in Th1 cells (IFNγ+CD4+TCRβ+) **(Fig. S4C)**, which augment immunity against intracellular pathogens such as STm (Santos et al., 2009). SFB also induces the fucosylation of epithelial cells via type 3 innate lymphoid cell (ILC3) signaling (Goto et al., 2014). We detected increased levels of UEA-1+Epcam+ fucosylated ileal epithelial cells in *Nav1.8-Cre/DTA* mice compared to control littermates **(Fig. S4D)**, however, we did not detect differences in the proportions of Lin-Thy1+ ILC3 populations (NKp46+, CD4+RORγt+, CD4-RORγt+) in *Nav1.8-Cre/DTA* mice compared to control littermates **(Fig. S4E)**. STm has been shown to activate sympathetic neurons that signal to muscularis macrophages (Gabanyi et al., 2016). We did not observe differences in the proportions of muscularis macrophages in nociceptor-ablated mice **(Fig. S4F)**.

Taken together, these data indicate that nociceptors regulate microbial homeostasis in the ileum, in particular SFB levels, which sets the threshold for STm colonization. However, major immune populations are not dysregulated in nociceptor-ablated mice. Transplantation of SFB into nociceptor-ablated mice enhanced host protection by preventing early STm colonization of the ileum and reducing PP invasion and dissemination.

### Nociceptor-mediated host protection is not through maintenance of epithelial defenses or GI motility

Given that STm interacts closely with epithelial cells and immune cells, we next tested whether barrier defenses were dysregulated in nociceptor-ablated mice. We analyzed epithelial permeability, proliferation, and antimicrobial peptide expression since these factors could alter gut microbiota composition and resistance to STm. We did not detect differences in intestinal permeability between *Nav1.8-Cre/DTA* mice and control littermates, as assessed by translocation of fluorescent dye from intestinal lumen into circulation **(Fig. S3A)**. We also did not detect differences in intestinal epithelial proliferation (Epcam+Ki67+ cells) between *Nav1.8-Cre/DTA* mice and control littermates **(Fig. S3B)**. To determine whether barrier-associated functions were perturbed at homeostasis, we performed RNA-seq transcriptional analysis on isolated ileal epithelial cells from *Nav1.8-Cre/DTA* mice compared to control littermates. Few differentially expressed genes were uncovered overall (>2-fold, p<0.05) **(Fig. S3C)**. Tight junction and antimicrobial peptide transcripts did not differ between *Nav1.8-Cre/DTA* and control samples **(Fig. S3D-E)**.

Intraepithelial lymphocytes are some of the first immune cells to encounter STm and have been shown to play a role in controlling early stages of infection (Ismail et al., 2011). We analyzed intraepithelial lymphocyte populations in *Nav1.8-Cre/DTA* mice, but detected no major differences in TCRαβ+ cells (CD4+ and CD8+) and TCRγδ+ cells **(Fig. S3F)**. We also performed subset analysis of TCRγδ+ cells, and did not detect differences between Vγ1+, Vγ2+, Vγ7+, or Vδ4+ subsets from *Nav1.8-Cre/DTA* and control littermates **(Fig. S3G)**. Therefore, nociceptor regulation of microbial homeostasis and host defense likely does not occur through intestinal epithelial integrity or intraepithelial lymphocytes.

We next hypothesized that nociceptors could modulate STm host defense by regulating GI transit. The role of gut-extrinsic sensory neurons, specifically nociceptors, in controlling GI motility has not been thoroughly investigated (Bartho et al., 2008). We assessed whether gut motility was impacted in nociceptor-ablated mice using assays to examine total GI transit and colonic transit. Nociceptor-ablated mice (*Nav1.8-Cre/DTA* mice, subcutaneous RTX-injected mice, and intrathecal RTX-injected mice) displayed significantly slower total GI transit and colonic transit compared to control mice **(Fig. S5A-B)**.

Given that nociceptor ablation led to slower GI transit, we next tested whether accelerating GI transit could enhance protection against STm infection in nociceptor-ablated mice. We administered vehicle and RTX-treated mice with an osmotic laxative (Miralax) in their drinking water, which led to faster total GI transit times in these mice compared to control mice given regular drinking water **(Fig. S5C)**. When Miralax was administered during STm infection to accelerate GI transit, we observed an unexpected increase in STm burdens in the ileums and spleens of mice given Miralax compared to water controls **(Fig. S5D)**. To examine whether restoring motility prior to infection could prevent STm infection, we administered Miralax to mice 5 days before infection and returned mice to regular water during infection **(Fig. S5E)**. We observed a similar increase in STm levels in the ileum and spleen at 5 dpi when Miralax was given before infection **(Fig. S5E)**. As an alternative strategy to increase intestinal motility, we gavaged wildtype mice with carbachol, a cholinergic agonist, which accelerated GI transit time **(Fig. S5F)**. Daily gavages of carbachol during STm infection led to increased STm levels in the Peyer’s patch and spleen at 5 dpi **(Fig. S5G)**.

These data indicate that while nociceptor neurons play a significant role in regulating gut motility, this role is likely dissociated from neural mechanisms of host protection against STm infection.

### Nociceptors regulate GP2+ Microfold cells in the Peyer’s patches that mediate *Salmonella* invasion

Several enteric pathogens, including STm, invade host tissues by penetrating ileal Peyer’s patches (PP) (Carter and Collins, 1974; Rivera-Chavez et al., 2016), and exploit M cells within the PP as a major cellular entry point (Jepson and Clark, 2001; Jones et al., 1994). M cells are antigen-sampling epithelial cells in the follicle-associated epithelium (FAE) that have short microvilli and a thin glycocalyx, making them vulnerable to enteroinvasive pathogens (Mabbott et al., 2013; Miller et al., 2007; Ohno, 2016; Williams and Owen, 2015). Previous studies have shown that sensory nerves innervate the PP dome (Chiocchetti et al., 2008; Vulchanova et al., 2007), and that prions in the GI tract enter through M cells to infect PP-innervating neurons (Takakura et al., 2011).

We asked whether neurons could interact with M cells in the Peyer’s patch and thereby regulate STm invasion. Whole-mount immunostaining of ileal PP FAE demonstrated that Tuj1+ nerve fibers were juxtaposed to M cells expressing glycoprotein 2 (GP2) (Fig. 5A and **Movie S1)**. GP2 is a GPI-anchored protein expressed on mature M cells (Kobayashi et al., 2013). STm uses a type-1 fimbrial adhesin, FimH, on its outer membrane to bind to GP2 to initiate bacterial invasion into M cells (Hase et al., 2009; Schierack et al., 2015). To investigate whether nociceptor-mediated host defense depended on STm binding to M cells for PP invasion, *Nav1.8-Cre/DTA* mice and control littermates were infected with a wildtype (WT) STm strain or an isogenic mutant strain lacking FimH (*∆FimH*). As previously observed, *Nav1.8-Cre/DTA* mice infected with WT STm had significantly greater bacterial invasion into PPs compared to control littermates (p=0.003). By contrast, almost none of the mutant ∆*FimH* STm were able to invade the PP of either *Nav1.8-Cre/DTA* mice or control mice (Fig. 5B, left). The levels of WT and ∆*FimH* STm colonizing the ileum were greater in *Nav1.8-Cre/DTA* compared to control mice (Fig. 5B, right), but reached similar levels in the cecum **(Fig. S6A)**, indicating that nociceptors regulate colonization resistance in the ileum and ileal PP invasion. Of note, the number of PPs, follicle number, or follicle area did not differ between *Nav1.8-Cre/DTA* and control littermates **(Fig. S6B-D)**. Thus, nociceptor regulation of host defense against STm depends on PP invasion, which requires an interaction between *Salmonella-* derived *FimH* and host GP2+ cells.

**Figure 5.**
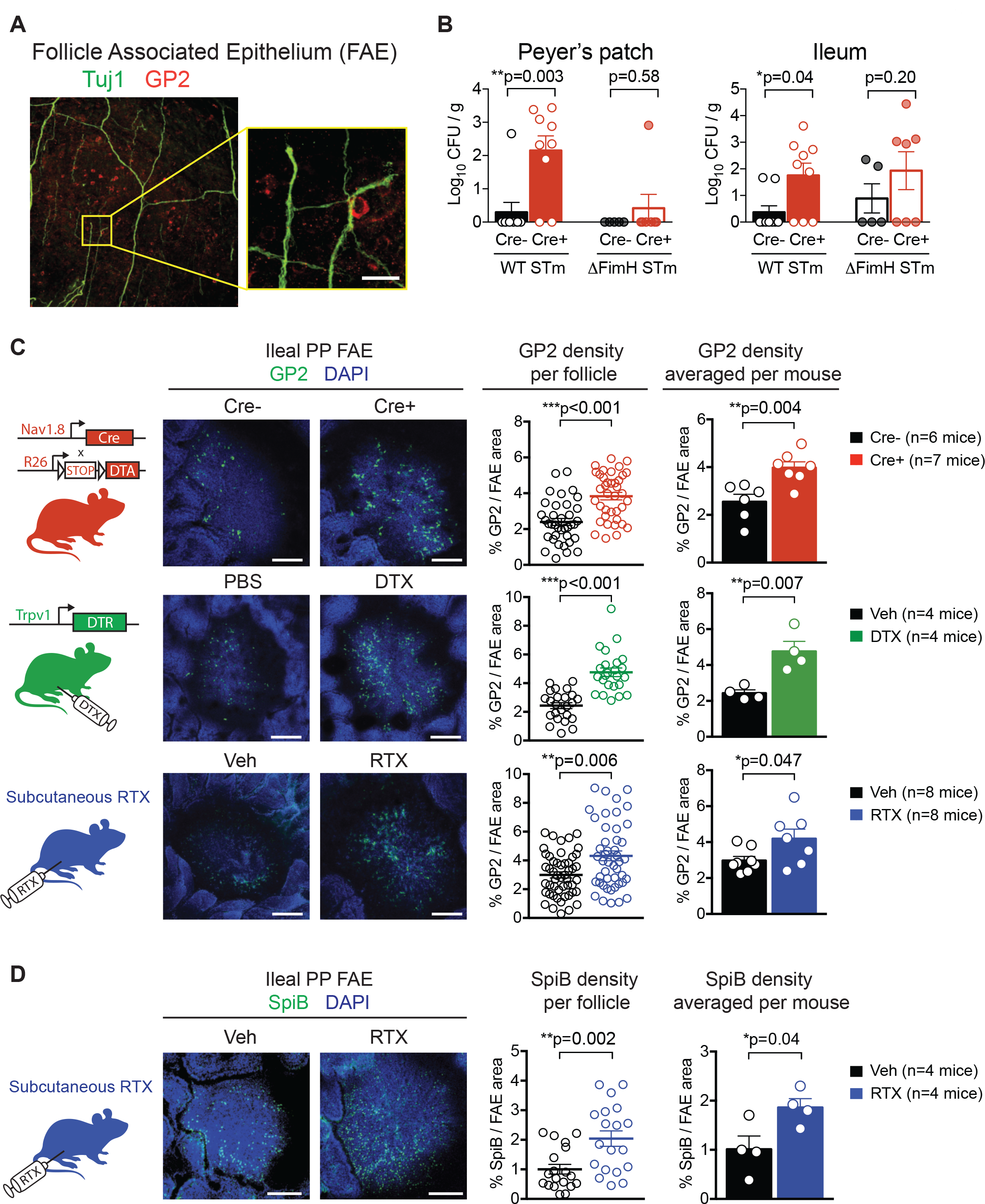
Nociceptor neurons regulate levels of Microfold (M) cells within the Peyer’s patch. **(A)** Whole-mount image of Peyer’s patch (PP) follicle associated epithelium (FAE) stained for Tuj1+ nerve fibers and glycoprotein 2 (GP2+) M cells. Scale bar inset, 25um. **(B)** *Nav1.8-Cre/DTA* mice and control littermates were infected with 0.8 × 10^7^ CFU of wildtype STm or 0.9 × 10^7^ CFU mutant Δ*FimH* STm and sacrificed at 1 dpi to analyze STm tissue burden. Data are pooled from 3 experiments with Cre-mice infected with WT STm (n=9), Cre-infected with Δ*FimH* STm (n=5), Cre+ mice infected with WT STm (n=9), and Cre+ infected with Δ*FimH* STm (n=7). **(C)** Quantification of GP2 density in ileal PP FAE from uninfected *Nav1.8-Cre/DTA*, *Trpv1-DTR*, and subcutaneous RTX-treated mice. Data are represented as GP2 density per follicle (left), or GP2 density of 3-6 follicles averaged per mouse (right). Data are pooled from 2 experiments of *Nav1.8-Cre/DTA* mice (n=6 Cre- and n=7 Cre+), *Trpv1-DTR* mice (n=4 vehicle and n=4 DTX), and RTX-treated mice (n=8 vehicle and n=8 RTX). Scale bar, 100um. **(D)** Quantification of SpiB density in ileal PP FAE from uninfected subcutaneous RTX-treated mice. Data from vehicle (n=4) and RTX (n=4) mice. Scale bar, 100um. Statistical analysis: **(B)** Non-parametric one-way ANOVA and Dunn’s post-tests for CFU analysis. **(C-D)** Unpaired t-tests for GP2 and SpiB analysis. *p<0.05, **p<0.01, ***p<0.001. Error bars are mean ± SEM. See also Figure S5 and S6.

Given that we observed increased STm burden in the PPs of nociceptor-ablated mice (Fig. 1B-D), and that M cells represent a bottleneck for STm invasion (Jepson and Clark, 2001), we hypothesized that nociceptors may regulate homeostasis of M cells and resulting invasion of STm into PPs. We analyzed the abundance of M cells in ileal PPs from nociceptor-ablated mice, including *Nav1.8-Cre/DTA, Trpv1-DTR*, and RTX-treated mice (Fig. 5C). We found significantly increased GP2+ M cell density in individual ileal PP FAEs, as well as increased GP2+ density averaged across mice in *Nav1.8-Cre/DTA* (p=0.004), *Trpv1-DTR* (p=0.007), and RTX-treated (p=0.047) mice compared to their controls (Fig. 5C). Upon infection, STm rapidly induces more M cells (Savidge et al., 1991; Tahoun et al., 2012). At 5 dpi, we observed higher levels of GP2+ M cells in RTX-treated mice compared to vehicle-treated mice **(Fig. S7A)**. We also analyzed levels of M cells expressing SpiB+, a second marker for M cells (Fig. 5D). SpiB is a transcription factor involved in early differentiation of M cells, and thus marks a large proportion of M cells (Kanaya et al., 2012; Sato et al., 2012). SpiB density per follicle and averaged across individual mice were also increased in RTX-treated mice compared to vehicle-treated mice (Fig. 5D).

Together, these data indicated that nociceptors suppress the density of M cells in ileal PPs and the absence of neurons led to significantly more M cells in the FAE at homeostasis and after STm infection.

### Nociceptor suppression of M cells protects against *Salmonella* infection

Given that M cells are primary entry points used by STm for invasion, nociceptive control of M cell density could play a critical role in restricting STm infection. To test this hypothesis, we targeted M cells in nociceptor-ablated animals by injecting vehicle or RTX-treated mice with neutralizing antibodies against the receptor activator of nuclear factor kappa-B ligand (RANKL). RANKL binds to its receptor RANK on epithelial cells to mediate the differentiation and maturation of M cells, and neutralizing anti-RANKL treatment was shown to lead to their transient depletion (Knoop et al., 2009; Nagashima et al., 2017). We found that anti-RANKL administration for 5 days led to a drastic reduction of GP2+ M cells in ileal PP FAE of vehicle and RTX-treated mice (Fig. 6A). Upon infection, isotype-treated RTX mice had significantly higher STm load in their spleens and livers compared to isotype-treated vehicle mice (Fig. 6B). By contrast, anti-RANKL-treatment in RTX-treated mice led to a drastic reduction of STm dissemination and weight loss compared to isotype-treated mice, thereby effectively restoring host defense in the nociceptor-ablated mice (Fig. 6B).

**Figure 6.**
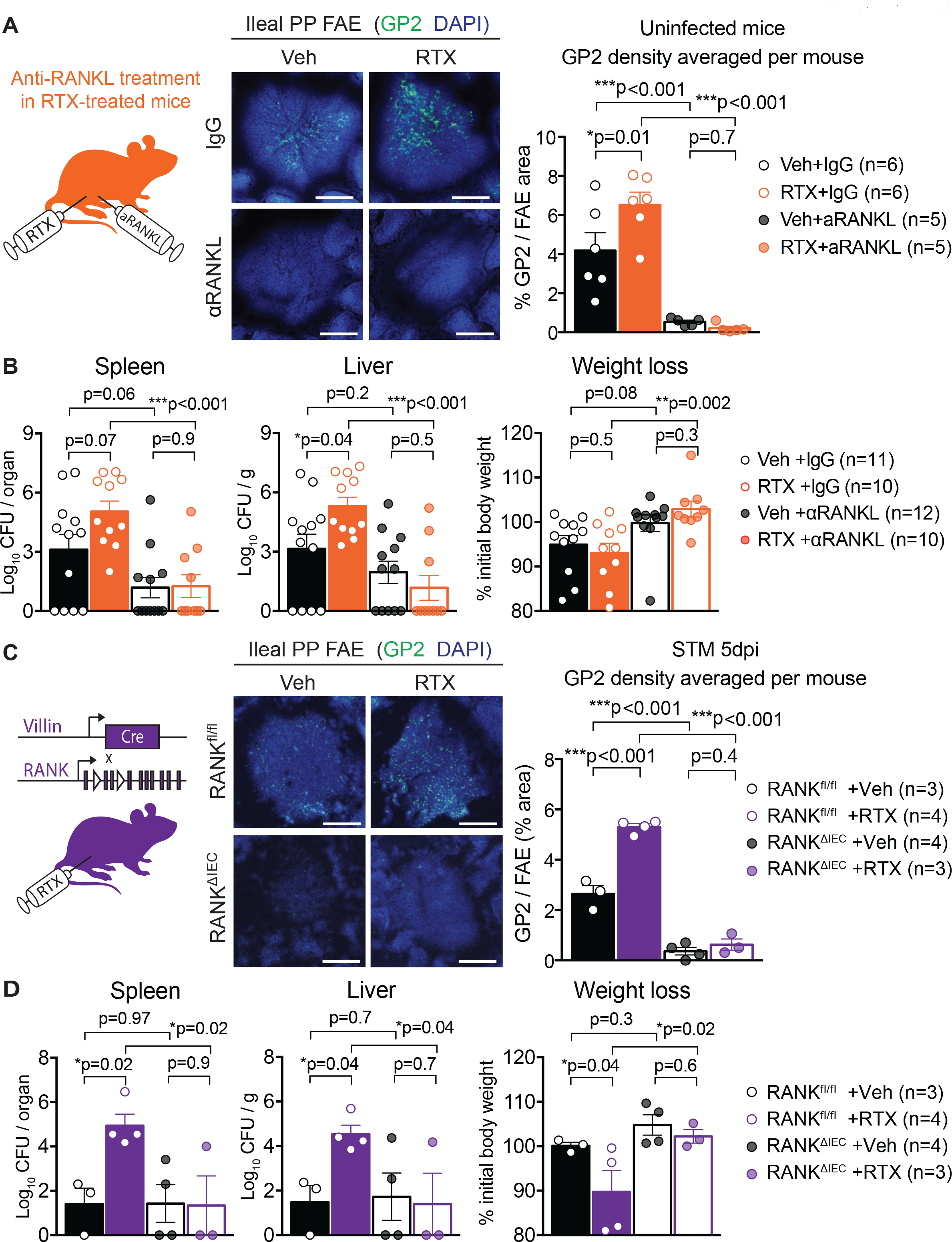
Blockade of RANK-RANKL signaling restores protection against *Salmonella* infection in nociceptor-ablated mice. **(A)** M cells were depleted in vehicle and RTX-treated mice by injections of anti-RANKL or IgG isotype control (250ug/mouse i.p. every 2 days starting 5 days prior to STm oral infection). Quantification of GP2 density in ileal PP FAE from 2 experiments of uninfected vehicle+IgG (n=6), RTX+IgG (n=6), vehicle+anti-RANKL (n=5), and RTX+anti-RANKL (n=5) treated mice. Data are represented as GP2 density per follicle (left), or GP2 density of 3-6 follicles averaged per mouse (right). Scale bar, 100um. **(B)** Following 5 days of pre-treatment with anti-RANKL or IgG control, vehicle and RTX-treated mice were orally infected with 0.7-1.9 × 10^7^ CFU STm, and sacrificed at 5 dpi to examine STm tissue burden and weight loss. Data are pooled from 3 experiments with vehicle+IgG (n=11), RTX+IgG (n=10), vehicle+anti-RANKL (n=12), and RTX+anti-RANKL (n=10) treated mice. **(C)** *Villin-Cre*^+/−^ mice were crossed with *RANK*^*fl/fl*^ mice to generate M-cell sufficient mice (RANK^fl/fl^) or mice with conditional epithelial cell depletion of RANK that were depleted of M cells (RANK^ΔIEC^). Mice were injected s.c. with RTX to deplete TRPV1+ neurons, then orally infected with 0.8 × 10^7^ CFU STm. Quantification of GP2 density in ileal PP FAE from RANK^fl/fl^+Veh (n=3), RANK^fl/fl^+RTX (n=4), RANK^ΔIEC^+Veh (n=4), and RANK^ΔIEC^+RTX (n=3) treated mice at STM 5dpi. Data are represented as GP2 density per follicle (left), or GP2 density of 3-6 follicles averaged per mouse (right). Scale bar, 100um. **(D)** Vehicle and RTX-treated Villin-Cre;RANK^fl/fl^ mice were sacrificed at 5 dpi to examine STm tissue burden and weight loss. Data are from RANK^fl/fl^+Veh (n=3), RANK^fl/fl^+RTX (n=4), RANK^ΔIEC^+Veh (n=4), and RANK^ΔIEC^+RTX (n=3) mice. Statistical analysis: **(A,C)** One-way ANOVA and Sidak post-tests for GP2 analysis. **(B,D)** Non-parametric one-way ANOVA and Dunn’s post-tests for CFU analysis. One-way ANOVA and Fisher’s post-test for weight loss analysis. *p<0.05, **p<0.01, ***p<0.001. Error bars are mean ± SEM. See also Figure S6.

We utilized a second M cell depletion strategy to assess their role in nociceptor-mediated host defense. We bred *Villin-Cre* mice with *RANK*^*fl/fl*^ mice to generate *Villin-Cre;RANK*^*fl/fl*^ mice (*RANK*^*∆IEC*^) that conditionally lacked RANK receptor signaling in gut epithelial lineage cells, thus resulting in M cell depletion (Rios et al., 2016). We treated *RANK*^*∆IEC*^ and control littermates (*RANK*^*fl/fl*^) with vehicle or RTX to eliminate TRPV1+ neurons. We confirmed that GP2+ M cells were absent in both vehicle and RTX-treated *RANK*^*∆IEC*^ mice by immunostaining of the FAE (Fig. 6C). Upon infection, STm loads in spleen and liver were significantly diminished when epithelial cells lacked RANK (Fig. 6D). Again, the differences between vehicle and RTX-treated mice in weight loss and STm dissemination were eliminated in the absence of M cells following infection (Fig. 6D). Taken together, these data indicate that nociceptor regulation of M cells plays a crucial role in STm host defense.

### Nociceptors do not regulate major PP cell populations or IgA levels

We next investigated how nociceptors may regulate M cell levels. We first analyzed PP epithelial proliferation by flow cytometry, but did not detect major differences in Epcam+Ki67+ cells before or after infection **(Fig. S6E)**. A prior study showed that the expression of RANKL by PP stromal cells mediates M cell differentiation (Nagashima et al., 2017). We performed qRT-PCR of PP tissues from vehicle and RTX-treated mice, but did not detect differences in transcript levels of either *RANKL* or osteoprotegerin (*OPG*), a negative regulator of RANKL signaling (Boyce and Xing, 2007) **(Fig. S6F)**. We also analyzed PP stromal cell populations, and found no difference in proportions of fibroblast reticular cells (FRCs), blood endothelial cells (BECs), or lymphatic endothelial cells (LECs) between vehicle and RTX-treated mice **(Fig. S6G)**. In addition to stromal cells, we analyzed whether other PP immune cell populations were regulated in nociceptor-ablated mice. Myeloid cells have been shown to be critical for phagocytosis and spread of STm from PP to mesenteric LNs (Vazquez-Torres et al., 1999). However, we did not detect major differences in F4/80+ macrophages and CD11b-dendritic cells (DCs), although we did see a slight increase in CD11b+CD11c+MHC-II+ DCs in RTX-treated mice **(Fig. S6H)**. There were also no differences in CD3+ T cells (CD4+, CD4-) **(Fig. S6I)** or B220+ B cells **(Fig. S6J)**. Additional analysis of IgA+ plasma B cells and GL7+ germinal center B cells did not reveal differences in the proportions of these cell subsets **(Fig. S6K)**. We also measured IgA levels in the feces, which have a major role in modulating STm agglutination and reducing invasion (Bioley et al., 2017), but found no differences between vehicle and RTX-treated mice, nor *Nav1.8-Cre/DTA* mice and control littermates **(Fig. S7B)**. Taken together, these data indicate that while PP M cell numbers are significantly different in the absence of nociceptors, the overall populations of PP immune and stromal cell-types are not grossly affected by nociceptors, nor IgA levels in the feces.

### Nociceptors directly detect STm and release the neuropeptide CGRP

Nociceptors can directly sense microbial products that are potentially harmful to the host (e.g. pore-forming toxins, lipopolysaccharides, flagellin, formyl peptides) (Blake et al., 2018; Chiu et al., 2013; Xu et al., 2015). Furthermore, nociceptors release neuropeptides that act on immune cells and other non-neural cell-types to modulate host inflammatory responses (Baral et al., 2018; Kashem et al., 2015; Maruyama et al., 2017; Talbot et al., 2015). We hypothesized that nociceptors could actively mediate host detection of STm and subsequently release neuropeptides that play a role in M cell homeostasis.

To test this hypothesis, we stimulated DRG neuronal cultures with medium or STm, and measured calcium influx by Fura-2 ratiometric imaging (Fig. 7A and **Fig. S7C)**. A subset of DRG neurons robustly responded to STm during incubation, which included TRPV1+ neurons as determined by their subsequent response to capsaicin. Calcium influx in neurons leads to SNARE-dependent release of neuropeptides, including CGRP, by peptidergic nociceptors (Sudhof, 2012). We found that STm induced DRG neuronal release of CGRP in a dose-dependent manner (Fig. 7B). Both heat-inactivated STm and supernatant from live STm cultures induced CGRP release. Pre-treatment of neurons with botulinum neurotoxin A (BoNT/A), which blocks SNARE-dependent vesicle fusion (Yang and Chiu, 2017), inhibited STm supernatant-induced CGRP release. As a positive control, capsaicin-induced activation of TRPV1+ neurons also stimulated CGRP release. Therefore, TRPV1+ DRG neurons are able to directly respond to STm by calcium influx and release of CGRP.

**Figure 7.**
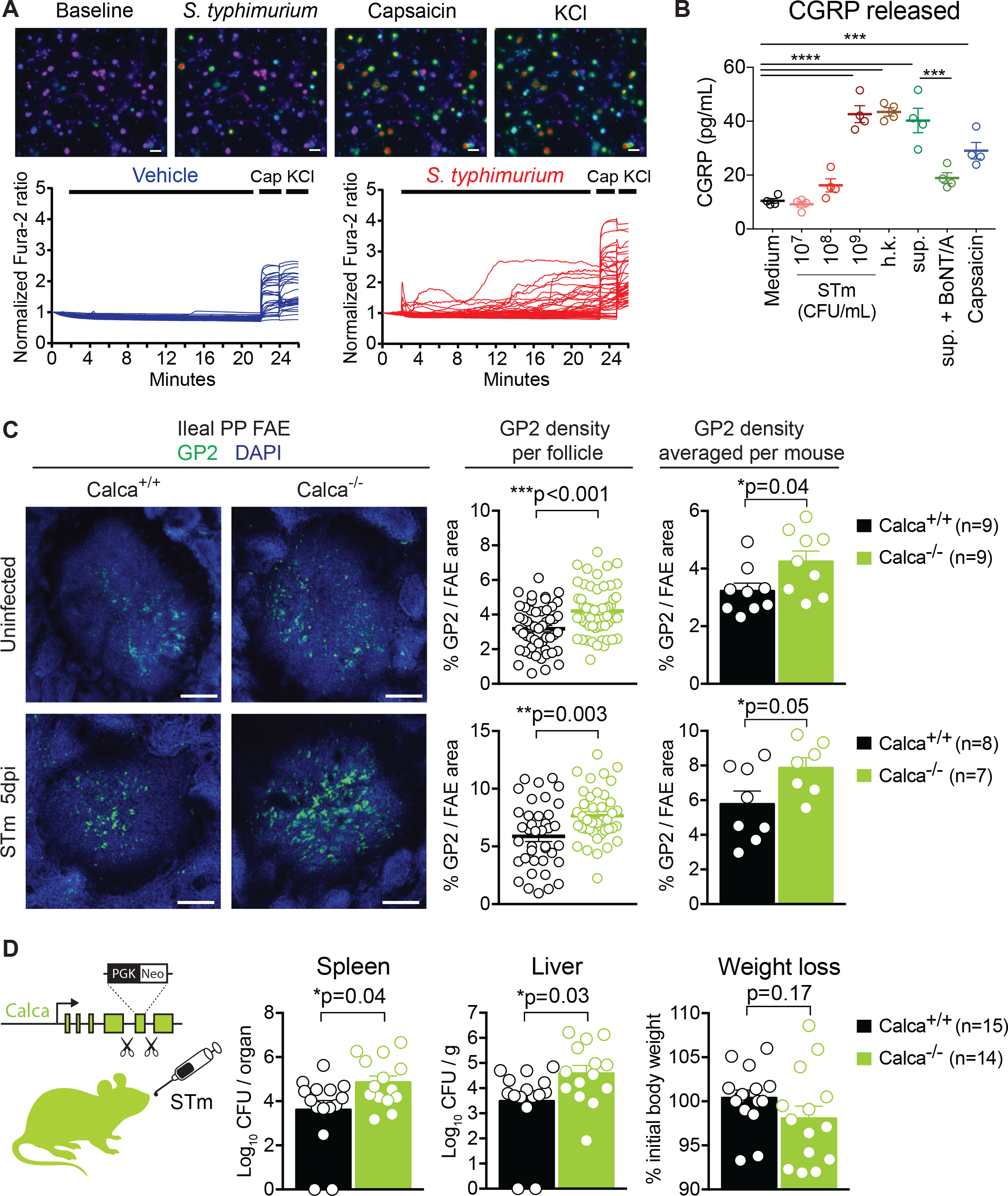
*Salmonella* induces DRG neuronal activation and release of CGRP, which can modulate M cells and *Salmonella* infection. **(A)** Representative calcium fields and traces of cultured DRG neurons stimulated with vehicle (Krebs-Ringer), STm (1 × 10^9^ CFU/ml), capsaicin (1uM Cap) and KCl (40mM) (n=5 plates/condition). Scale bar, 30um. **(B)** CGRP released from cultured DRG neurons stimulated with live, heat-killed, and supernatant cultures of STm. Supernatant-induced CGRP release is blocked by addition of Botulinum toxin (BoNT/A) (n=4 samples/condition). **(C)** Quantification of GP2 density in ileal PP FAE of *Calca*^+/+^ and *Calca*^−/−^ littermates. Data are represented as GP2 density per follicle (left), or GP2 density of 3-6 follicles averaged per mouse (right). Data are pooled from 2 experiments with uninfected *Calca*^+/+^ (n=9) and *Calca*^−/−^ (n=9) mice, and from *Calca*^+/+^ (n=8) and *Calca*^−/−^ (n=7) mice infected at STM 5dpi. Scale bar, 100um. **(D)** *Calca*^+/+^ (n=15) and *Calca*^−/−^ (n=14) mice were orally infected with STm and sacrificed at 5 dpi to examine STm dissemination and weight loss. Data are pooled from 3 experiments with *Calca*^+/+^ (n=15) and *Calca*^−/−^ (n=14) mice infected with 1.9-2.2 × 10^7^ CFU STm. Statistical analysis: **(B)** One-way ANOVA and Sidak post-tests for CGRP release. **(C)** Unpaired t-tests for GP2 analysis. **(D)** Mann-Whitney tests for CFU analysis. Unpaired t-test for weight loss analysis. *p<0.05, **p<0.01, ***p<0.001. Error bars are mean ± SEM. See also Figure S7.

### CGRPα modulates M cells and *Salmonella* host defense

The role of CGRP in enteric host defense against STm has not been previously investigated. CGRP is expressed as 2 isoforms, CGRPα (*Calca*) and CGRPβ (*Calcb*), which have similar functional activity, but are differentially distributed in gut extrinsic and intrinsic neurons (Fig. S1A) (Mulderry et al., 1988). We showed that our nociceptor targeting strategies eliminated extrinsic CGRP+ fibers entering the GI tract, while intrinsic CGRP+ (i.e. *Calcb*) neurons were left intact (Fig. 2C-E). Both *Calca* and *Calcb* transcripts were significantly reduced in the DRG of *Nav1.8-Cre/DTA* mice, whereas in myenteric plexus tissues, *Calca* was not detected and *Calcb* was unaffected compared to control littermates **(Fig. S7D)**. Given that we demonstrated that extrinsic DRG nociceptors mediated STm host defense (Fig. 2F), and that CGRPα (*Calca*) is expressed at high levels in DRG neurons (Evangelista, 2014), we next determined whether this isoform played a role in STm host defense. Using mice with targeted disruption of exon 5 of the *Calca* gene (*Calca*^−/−^ mice), we found that the abundance of GP2+ M cells was significantly higher in ileal PP FAE of *Calca*^−/−^ mice compared to *Calca*^+/+^ control littermates (Fig. 7C). We next examined mice that had targeted disruption of exon 3 and 4 of the *Calcb* gene (*Calcb*^−/−^ mice). By contrast with *Calca*^−/−^ mice, we found no difference in GP2+ cell density between *Calcb*^+/+^ and *Calcb*^−/−^ littermates **(Fig. S7E)**.

Given the effect of *Calca* in regulating M cell numbers at homeostasis, we next examined whether this isoform played a role in STm infection. After 5 dpi, *Calca*^−/−^ mice also showed significantly higher GP2+ M cell densities than *Calca*^+/+^ littermates (Fig. 7C). We tested the possibility that CGRP could directly inhibit the differentiation of M cells using epithelial organoids cultured from small intestinal crypts. Treatment of epithelial organoids with RANKL led to an upregulation of *Gp2* and *Spib* transcripts, which were not significantly affected by the addition of CGRP **(Fig. S7F)**. Finally, to examine whether CGRPα (*Calca*) could contribute to host defense during STm infection, we infected *Calca*^−/−^ mice with STm. Following oral infection, we found that *Calca*^−/−^ mice had significantly increased STm burden in the spleens (p=0.04) and livers (p=0.03) compared to *Calca*^+/+^ control littermates (Fig. 7D). Therefore, nociceptor neurons can directly respond to STm infection by releasing CGRP, and CGRPα regulates M cell numbers to improve host defense against STm.

## DISCUSSION

While nociceptor neurons are major sensors of disturbances in the GI tract and mediate pain, nausea, and gastrointestinal inflammation, their role in host defense against enteric pathogens has not been explored. In this study, we found that nociceptor neurons protected the host against GI infections of STm through two mechanisms: by regulating small intestinal microbiota composition to facilitate colonization resistance, and by suppressing Peyer’s patch M cell abundance to minimize pathogen invasion. These results uncovered a role for nociceptor neurons in regulating barrier defenses in the GI tract that has wider implications for the treatment of both inflammation and infection.

### Nociceptor regulation of the small intestinal microbiota

The gut microbiota is shaped at homeostasis by many factors in the host micro-environment, including factors produced by gut-resident epithelial and immune cells such as mucins, antimicrobial peptides, and immunoglobulins (Duerkop et al., 2009; Garrett et al., 2010). In this study, we demonstrate that the nervous system also plays an important role in shaping the gut microflora. Nociceptor neurons specifically regulated bacterial communities in the small intestine ileum, but not in the duodenum, colon or feces. Future work could determine how nociceptors modulate the intestinal microbiota. It is possible that nociceptors regulate bacterial communities indirectly through modulation of epithelial cell function (Sharkey et al., 2018; Yoo and Mazmanian, 2017). Epithelial cells, including goblet cells and enterochromaffin cells, possess neurotransmitter and neuropeptide receptors (Knoop and Newberry, 2018; Snoek et al., 2010; Vergnolle and Cirillo, 2018). Alternatively, there is evidence that neurotransmitters, including catecholamines, released into the gut lumen can directly modify bacterial growth and production of virulence factors (Hughes and Sperandio, 2008). CGRP and substance P, which are neuropeptides produced by nociceptors, have been shown to have direct antimicrobial effects on enteric, skin, and oral pathobionts (Augustyniak et al., 2012; El Karim et al., 2008). Examining the mechanisms underlying neural modulation of the microbiota could yield novel therapeutic targets that impact gut homeostasis and host defense.

We found that nociceptors maintained SFB levels in the ileum mucosa and lumen, which contributed to host protection against STm infection. SFB has been shown to limit pathogen colonization by *Salmonella enteriditis, Escherichia coli*, and *Citrobacter rodentium* (Garland et al., 1982; Heczko et al., 2000; Ivanov et al., 2009). Here we show that SFB can also protect against *Salmonella typhimurium* infection. Restoring SFB levels in nociceptor-ablated mice significantly reduced STm colonization of the ileum and curtailed downstream parameters of infection, including invasion and dissemination. One possibility is that SFB regulates mucosal immunity to hinder STm survival and/or invasion. However, we did not detect major differences in the abundance of major immune cell types between *Nav1.8-Cre/DTA* mice and control littermates before or after infection. Another possibility is that SFB occupies the small intestinal niche where STm invades and competes physically with STm for binding spots on ileal epithelial cells or metabolically for small intestine nutrients (Garland et al., 1982). SFB could also produce mediators that shape the bacterial community within the ileum that then sets the threshold for STm invasion. We found that the decrease in SFB led to increased levels of other bacterial members (e.g. *Prevotella, Ruminococcus*, etc.) in nociceptor-ablated mice. These changes led to an increase in the *Bacteroidetes*/*Firmicutes* ratio, which previous research has linked to an increased susceptibility to intestinal inflammatory conditions, such as colitis (Walters et al., 2014).

### Nociceptor regulation of gastrointestinal motility

We also found that extrinsic nociceptors regulated GI motility at homeostasis. While neural control of GI motility by gut-extrinsic neurons, in particular the sympathetic and parasympathetic nervous systems have been well described, data supporting a role for extrinsic sensory neurons in regulating GI motility is relatively weak (Bartho et al., 2008). Our work demonstrated that nociceptor-ablated mice had disturbances in GI motility with slower total and colonic GI transit compared to control mice. This could occur through nociceptor release of bioactive neuropeptides from their peripheral nerve terminals that act on enteric neurons or smooth muscle, or through activation of spinal or brain circuits that signal back to the intestine via efferent outflow (i.e. sympathetic and parasympathetic neurons). We found that nociceptor regulation of GI motility was separate from their role in host defense since accelerating GI transit by using an osmotic laxative (Miralax) or a cholinergic agonist (carbachol) led to worsened outcomes of STm infection. This is contrary to the long-held hypothesis that accelerating peristalsis, along with hypersecretion of water and electrolytes (i.e. “infectious diarrhea”) should aid in the clearance of acute GI microbial infections (Hodges and Gill, 2010). There are several reasons why accelerating GI transit may not be protective. Facilitating GI transit using osmotic laxatives significantly depletes the mucus layer (Tropini et al., 2018) and greatly perturbs the commensal microbiota (Gorkiewicz et al., 2013), which could lead to dysbiosis and promote pathogen expansion. The effects of accelerating GI motility on infection outcomes may be pathogen-specific since contrary to what we observed with STm infections, Miralax treatment protected against colonization of *Citrobacter rodentium* in the colon (Tsai et al., 2017). Future studies are necessary to explore the relationship between different neural subsets (e.g. sensory, autonomic), GI motility, and infection given that specific neuro-immune mechanisms could contribute to disparate phenotypes.

### Nociceptor regulation of Peyer’s patch M cells

We found that nociceptors suppressed the abundance of M cells in ileal PPs at homeostasis and during STm infection. M cells are important sentinel cells that initiate mucosal immunity by transporting luminal antigens to underlying professional antigen-presenting phagocytes (Miller et al., 2007; Ohno, 2016; Williams and Owen, 2015). Enteroinvasive pathogens (e.g. *Salmonella, Shigella, Yersinia*) exploit M cells by using them as entry points to invade the small intestine (Jepson and Clark, 2001; Jones et al., 1994; Sansonetti and Phalipon, 1999). Additionally, STm itself can significantly enhance the numbers of M cells upon infection by mediating transdifferentation of these cells (Tahoun et al., 2012). In our study, we found that nociceptor modulation of M cells created a bottleneck for STm invasion into PPs and restricted systemic dissemination.

We found that nociceptor regulation of M cell abundance and STm infection occurs in part via CGRP. DRG nociceptors are directly activated by STm and respond by releasing CGRP. We found that mice lacking CGRPα, but not CGRPβ, had increased M cell density in ileal PPs and greater STm dissemination compared with control littermates. Nociceptors can release several soluble mediators during activation including substance P (SP), vasoactive intestinal peptide (VIP), and somatostatin (SST), thus CGRPα could be one of many mediators through which nociceptors confer host protection against enteric pathogens. How CGRP regulates M cell levels remains unclear. CGRP acts through its receptor RAMP1/CALCRL, which is highly expressed by both immune cells (Assas et al., 2014) and goblet cells (Haber et al., 2017). CGRP can increase goblet cell production and secretion of mucins (Ichikawa et al., 2000; Plaisancie et al., 1998), which may further impact microbial composition in the small intestine and microbial proximity to epithelial cells (Donaldson et al., 2016b). It is also possible that CGRP could regulate M cells via modulation of RANKL signaling, which has been shown to be critical for the differentiation and maturation of M cells (Knoop et al., 2009; Nagashima et al., 2017). Studies in osteoclasts showed that CGRP suppressed RANKL signaling to affect bone remodeling (Maruyama et al., 2017; Takahashi et al., 2016). Although we did not detect major differences in RANKL expression in PP of nociceptor-ablated mice, we nevertheless found that blockade of RANKL signaling rescued the host defense phenotypes. Hence, neural control of M cell differentiation could intersect with multiple cellular and molecular signaling pathways during gut homeostasis that require further investigation.

It may make physiological sense that gut-innervating nociceptor neurons, whose fundamental role is to protect mammals from danger, would also limit pathogen entry points by regulating M cell numbers. This neural control could occur at different levels during gut homeostasis. Detection of noxious stimuli, such as mechanical injury or diet-derived irritants, could lead to neural suppression of M cell numbers to protect the gut from future exposure to pathogens or toxins, and this process could further accelerate when a pathogen invades the gut.

There is growing evidence that nociceptor neurons, in addition to detecting classic noxious stimuli (e.g. noxious heat, ATP, reactive chemicals) (Basbaum et al., 2009; Woolf and Ma, 2007), can directly sense bacterial pathogens (Yang and Chiu, 2017), and initiate neuro-inflammatory processes (Chavan et al., 2017; Talbot et al., 2016). Nociceptors also crosstalk with immune cells to modulate host defense against *S. aureus, S. pyogenes*, and *C. albicans* (Baral et al., 2018; Kashem et al., 2015; Pinho-Ribeiro et al., 2018). In this study, we find that nociceptor neurons directly detect *Salmonella typhimurium*. Some candidate molecules from STm that could activate neurons include flagellin, as TLR5 has been found to be expressed by sensory neurons (Xu et al., 2015) and LPS, which has been found to directly activate the nociceptive ion channel TRPA1 (Meseguer et al., 2014). Future experiments will determine the exact mechanisms by which neurons sense *Salmonella* and other enteric pathogens.

Recent work has shown that several neuronal sub-types play a key role in neuro-immune signaling in the GI tract during inflammation and host defense. Cholinergic enteric neurons modulate type 2 innate lymphoid cells (ILC2s) via the neuropeptide neuromedin U (NMU) to significantly impact helminth infections (Cardoso et al., 2017; Klose et al., 2017). Gut-innervating sympathetic neurons polarize muscularis macrophages through β_2_AR+ adrenergic receptors toward a tissue-protective phenotype at homeostasis and after *Salmonella* infection (Gabanyi et al., 2016). Muscularis macrophages also cross-talk with M-CSF-releasing enteric neurons by releasing BMP7 to modulate GI motility (Muller et al., 2014), and self-maintaining gut macrophages perform niche-specific functions to promote intestinal homeostasis, including motility (De Schepper et al., 2018).

While neuronal regulation of inflammation in the gut has been described by sympathetic neurons and intrinsic enteric neurons (Cervi et al., 2014; Sharkey and Savidge, 2014), less is known about the extrinsic sensory neuronal contribution. In the gut, extrinsic nociceptors mediate visceral pain sensation and nausea/vomiting reflexes, which are prominent symptoms of enteric pathogen infections (e.g. *E. coli, Salmonella, Vibrio*). Our study expands the role of gut-innervating nociceptors by showing that they modulate several layers of host GI physiology, including regulating small intestine microbial communities, gut motility, and Peyer’s patch M cell numbers to impact host defense. Uncovering the mechanisms underlying neural modulation of the immune system and microbiota to impact gut homeostasis and host defense remain rich areas for future exploration. These studies are potentially useful for developing host-directed treatments that target distinct GI infections, and could contribute to alleviating the massive global health burden posed by emerging drug-resistant enteric pathogens.

## ACKNOWLEDGEMENTS

We thank Alice Cui, Marie Siwicki, Trevor Krolak, Sebastien Sannajust, Wanyin Tao, Fabian Chavez Rivera, Jonathan Chang, Pallab Ghosh, Adrianus Van Der Velden, Nicole Lee, Joseph Borrell, Chad Araneo, Alos Diallo, and Paula Montero Llopis for helpful advice and technical support. We thank Neil Mabbott, David Donaldson, Muriel Larauche, Wendy Garrett, Stephen Liberles, Gerald Pier, Dennis Kasper, Christophe Benoist, Diane Mathis, Nissan Yissachar, Michael Carroll, and Darren Higgins for insightful discussions. We thank Evegeni Sokurenko (University of Washington) for generously providing the *ΔFimH* SL1344 strain. We thank Pablo Pereira (Institut Pasteur) for generously providing the Vγ7 antibody. We thank Darren Higgins, Andreas Baumler, Sebastian Winter, and Claire Bryant for other bacterial strains not used in this study. We thank Matthew Gudgeon and Tammy Hshieh for support. This work was generously supported by funding from National Institutes of Health (NIH) under NCCIH DP2AT009499 (I.M.C.), NIH/NIAID K22AI114810 (I.M.C.), the Harvard Digestive Disease Center (I.M.C. funded by NIH P30 DK348345), Chan-Zuckerberg Initiative (I.M.C.), Harvard Stem Cell Institute (I.M.C.), NIH/NIDDK K08 DK110532 (M.R.), NIH K08 AI108690 (N.K.S.), and Whitehead Scholar and Translating Duke Health Scholar (N.K.S.).

## AUTHOR CONTRIBUTIONS

Conceptualization and design of project – N.Y.L. and I.M.C.; Mouse infection studies – N.Y.L., P.M., D.E.P., A.M., H.S., D.P.; Imaging analysis – N.Y.L., F.A.P-R., M.A.M., P.M., D.E.P., K.G., K.N.S., S.S.; Microbiota and SFB analysis – N.Y.L., D.P., J.R.H., D.A., A.M., D.E.P.; Gut motility – N.Y.L., P.M., H.S., Z.C.; Flow cytometry and immune cell analysis – N.Y.L., P.B., A.M.; Neuronal analysis – F.A.P-R., M.A.M., P.M., M.R., S.S.; Data processing and plotting - N.Y.L., M.A.M., P.M., F.A.P.-R., D.P., P.B., D.E.P., A.M., H.S., K.N.S., S.S., Z.C., K.G.; Provision of key resources and funding – M.N.S., M.R., V.K.K., C.W., A.W., J.R.H., N.K.S., R.N., D.A.; Writing of the original draft – N.Y.L. and I.M.C.; Critical editing of manuscript – N.Y.L., M.A.M, A.J., I.M.C.

## METHODS

### Mice

C57BL/6 mice were purchased from Jackson Laboratories (Bar Harbor, ME) or Taconic Biosciences (Rensselaer, NY). *B6.Rosa26-stop(flox)-DTA* (Voehringer et al., 2008) and Ai14 strain *B6.Rosa26-stop(flox)-tdTomato* (Madisen et al., 2010) mice were purchased from Jackson Laboratories. *Nav1.8-Cre* (Abrahamsen et al., 2008) mice were provided by J. Wood (University College London). *Trpv1-DTR* (Pogorzala et al., 2013) mice were provided by M. Hoon (NIH). *Calca*^−/−^ (Oh-hashi et al., 2001) mice were provided by V. Kuchroo (Harvard Medical School). *Calcb*^−/−^ (Thompson et al., 2008) mice were provided by M. Rao (Columbia University). *Villin-Cre* (Madison et al., 2002) and *RANK*^*fl/fl*^ (Rios et al., 2016) mice were provided by N. Surana (Duke University). For Nav1.8-lineage neuron depletion experiments, *Nav1.8-Cre*^+/−^ mice were bred with *Rosa26-stop(flox)-DTA*^+/+^ mice to generate Nav1.8-lineage neuron-depleted (*Nav1.8-Cre*^+^/*DTA* or Cre+) mice and control (*Nav1.8-Cre*^−^/*DTA* or Cre-) littermates. For M cell depletion experiments, *Villin-Cre*^+/−^;*RANK*^*fl/fl*^ were bred to *Villin-Cre*^−/−^;*RANK*^*fl/fl*^ mice to generate mice with epithelial-specific depletion of RANK (*RANK*^*ΔIEC*^) or control (*RANK*^*fl/fl*^) littermates. *Nav1.8-Cre*^+/−^ mice were bred with *B6.Rosa26-stop(flox)-TdTomato*^+/+^ mice to generate *Nav1.8-Cre*^+^/*TdTomato*^+^ mice. For *Calca* and *Calcb* experiments, heterozygous mice were bred together to produce wildtype and knockout littermates. Mice were bred and housed in a specific pathogen-free animal facility at Harvard Medical School (HMS). Age-matched 7-to 14-week-old littermate mice of both genders were used for experiments. All animal experiments were approved by the HMS Institutional Animal Use and Care Committee.

### Bacterial strains and culture conditions

The wildtype *Salmonella typhimurium* (STm) strain SL1344 was provided by M. Starnbach (HMS). SL1344 *ΔFimH* was provided by E. Sokurenko (University of Washington). SL1344 wildtype strain was grown in LB broth, and *ΔFimH* mutant strain was grown in LB broth with 25 ug/ml kanamycin. For infection studies, bacteria were grown overnight for 16-18 hr at 37°C on a shaker, 250 r.p.m. in LB broth supplemented with antibiotics when appropriate. Bacteria were centrifuged at 5,000 r.p.m. for 5 min, washed twice, and resuspended in sterile PBS. The OD600 was measured to estimate bacterial density, and serial plating was performed on LB agar plates to quantify the infection dose by counting colony forming units (CFU) after an overnight incubation at 37°C.

### Bacterial infections and CFU determination

In murine typhoid-like infection models, mice were fasted overnight in clean cages with unlimited access to water. On the morning of infection, mice were gavaged with 100 ul of 3% sodium bicarbonate, followed 30 min later by 200 ul of bacteria resuspended in PBS at indicated doses. In some experiments, mice received an intraperitoneal injection with 200 ul of bacteria in PBS for systemic infections. To evaluate bacterial load, mice were euthanized with CO_2_ and tissues were rapidly dissected. Tissues were transferred into 2 ml eppendorf tubes containing ice-cold PBS and a 5 mm steel bead. Tissues were weighed and homogenized in a TissueLyzer (Qiagen) for 2-4 min at 30 Hz, then serially diluted in PBS for plating. Mesenteric lymph nodes, Peyer’s patches, spleens and livers were plated on LB agar. Ileum (distal 5cm without Peyer’s patches) and cecum were plated on LB agar with 50 ug/ml streptomycin. Freshly collected feces were plated on McConkey agar with 50 ug/ml streptomycin. Additionally, Peyer’s patches were treated with 100 uM gentamicin for 1 hr at 37°C to remove extracellular bacteria, washed twice with PBS before homogenizing and plating. Bacterial CFU were counted after an overnight incubation at 37°C.

### Genetic and chemical ablation of TRPV1 nociceptor neurons

For genetic ablation of TRPV1 neurons, *Trpv1-DTR* mice were treated with diphtheria toxin (DTX) as previously described (Pogorzala et al., 2013; Trankner et al., 2014). Mice were injected intraperitoneally (i.p.) with 200 ng of DTX (Sigma Aldrich) dissolved in 100 ul PBS daily for 21 days. Vehicle-treated mice were injected i.p. with 100 ul PBS. Mice were allowed to rest for at least 1 week before being used for experiments. For chemical ablation of TRPV1 neurons, 4-week-old C57BL/6 mice were treated with resiniferatoxin (RTX, from Sigma Aldrich) as previously described (Elekes et al., 2007; Szolcsanyi et al., 1990). For subcutaneous injections, mice were injected in the flank under isoflurane with three increasing doses of RTX (30, 70, and 100 ug/kg on consecutive days) dissolved in 2% DMSO/0.15% Tween-80/PBS. To target spinal TRPV1 neurons by intrathecal injections, 4-week-old C57BL/6 mice were injected with RTX (25 ng/mouse) dissolved in 10ul of 0.25% DMSO/0.02% Tween-80/0.05% ascorbic acid/PBS in the L5-L6 region using a 30G needle and Gastight syringe (Hamilton) under isoflurane on two consecutive days. For both subcutaneous and intrathecal injections, control mice were treated with vehicle alone. Mice were allowed to rest for 4 weeks before being used for experiments. Loss of TRPV1+ neurons in the DRG and vagal ganglia was confirmed by immunostaining, or by reduced thermal responses to noxious heat during hot plate tests (55°C).

### Immunostaining of extrinsic ganglia

For extrinsic ganglia, mice were perfused with PBS followed by 4% PFA/PBS. Lumbar DRG (L1-L6) and vagal ganglia were dissected, post-fixed for 2 hr in 4% PFA/PBS at 4°C, incubated overnight at 4°C in 30% sucrose/PBS, and embedded in OCT. Sections (14 um) were cut and stained with primary antibodies (rabbit anti-CGRP, Millipore, PC205L, 1:500; guinea-pig anti-TRPV1, Millipore, AB5566, 1:1,000; chicken anti-Neurofilament 200, Millipore, AB5539, 1:500). Sections were washed in PBS, then stained with secondary antibodies (donkey anti-rabbit DyLight 488, Abcam, 1:500; goat anti-guinea pig, Sigma, 1:500; anti-chicken Alexa 488, Life Technologies, 1:1000). Sections were mounted in VectaShield (Vector Labs) with DAPI, and imaged by an Olympus Fluoview 1000 confocal microscope with 10X magnification at HMS Microscopy Resources on the North Quad (MicRoN) Core. Maximum projection images obtained for each channel were exported for analysis. For quantification of extrinsic ganglia neurons, the % of TRPV1+, CGRP+, or NF200+ neurons out of the total Tuj1+ neurons was determined for each sample as an average of 3 fields per mouse. Investigators were blinded to genotypes and treatment groups.

### Immunostaining of myenteric plexus and mesentery

Immunostaining was performed as previously described (Rao et al., 2015). Ileums and attached mesentery were flushed with PBS and fixed in 10% formalin/PBS for 2 hr on ice. Longitudinal muscle layers and associated myenteric plexus were separated from the mucosa/submucosa. Tissues were washed in PBS, blocked in 5% normal donkey serum/1% Triton X-100/PBS for 2 hr at RT, then incubated overnight in primary antibodies at 4°C (rabbit anti-HuC/D, Abcam, ab184267 at 1:2000; goat anti-Calretinin, Swant, CG1 at 1:100; goat anti-nNOS, Abcam, ab1376 at 1:100; anti-rabbit CGRP, Millipore, AB5539 at 1:500). Following PBS rinses, samples were incubated in succession in secondary antibody solution for 2 hr at RT (donkey anti-rabbit IgH&L Alexa594, Abcam, ab150076; donkey anti-goat IgH&L Alexa488, Abcam, ab150129). Tissues were washed in 1% Triton X-100/PBS, then in PBS, and mounted in Aqua-Poly/Mount (Polysciences, 18606-20). Z-stacks (0.9um intervals) were acquired on a Nikon Ti inverted spinning disk confocal microscope with a 20X magnification using NIS Elements Acquisition Software AR 5.02 at HMS MicRoN Core. For quantification of myenteric plexus neuron populations, >800 neurons over >5 images were counted per mouse. The total HuD+ neurons, % of calretinin+ and nNOS+ neurons out of HuD+ neurons were calculated per mouse and plotted in GraphPad Prism. Investigators were blinded to genotypes and treatment groups.

### Immunostaining of Peyer’s patch M cells and neuron innervation

Immunostaining was performed as previously described (Kanaya et al., 2012; Kobayashi et al., 2013). For whole-mount staining, 1 cm segments of ileum tissues containing PPs were washed 5 times in 5% FBS/10mM DTT/0.05% Tween-20/1uM nicardipine/HBSS to remove mucus and maximize smooth muscle relaxation. Tissues were opened longitudinally, stretched out, and fixed for 1 hr on ice in Cytofix/Cytoperm buffer (BD Biosciences). Tissues were blocked in 10% normal goat serum or donkey serum/0.25% Triton X-100/PBS for 2-4 hr at 4°C, then incubated overnight at 4°C in primary antibodies in 0.1% Triton X-100/PBS (rat anti-GP2, MBL, D278-3 at 1:500; sheep anti-SpiB, R&D Systems, AF7204 at 1:50; rabbit anti-Tuj1, Abcam, ab18207 at 1:1000). Tissues were washed for 20 min, 5 times in PBS, then incubated in secondary antibodies in 0.1% Triton X-100/PBS for 2 hr at 4°C (goat anti-rat Alexa 488, Abcam, ab150157 at 1:500; donkey anti-rat Alexa 594, Jackson ImmunoResearch, 712-585-153 at 1:500; donkey anti-sheep Alexa 488, Abcam, ab150177 at 1:500; Hoechst 33342, Thermo 66249 at 1:10,000, final 2 ug/ml). Tissues were washed again, then mounted in VectaShield (Vector Labs). For M cell density in PP FAE, Z-stacks (3um interval) were acquired on an Olympus Fluoview 1000 confocal microscope with a 10X magnification; for FAE neuron innervation, Z-stacks (0.5um interval) and tiled images were acquired on a Nikon Ti inverted spinning disk confocal microscope with a 20X magnification at HMS MicRoN Core. To calculate the density of M cells, maximum projection images were compiled in ImageJ. The follicle area encircled by surrounding villi was traced using the lasso tool in Adobe Photoshop after setting a threshold for nuclei stained by Hoechst 33342. The follicle area region of interests (ROI) were imported into ImageJ, and an automated Otsu threshold for GP2 and SpiB channels was applied. The follicle area (um^2^) and % of follicle stained by GP2+ or SpiB+ cells was quantified in ImageJ and plotted in GraphPad Prism. Investigators were blinded to genotypes and treatment groups during imaging and analysis.

### Quantitative PCR

Whole DRG, vagal ganglia, myenteric plexus/longitudinal muscle from ileum and colon, PPs, and epithelial organoids were homogenized in Trizol reagent and stored at −80°C until RNA extraction. Total RNA was isolated by addition of chloroform to homogenates and centrifuged at 12,000 g for 15 min at 4°C. The aqueous phase was combined with equal parts of 70% ethanol and RNA was purified using an RNeasy Mini Kit (Qiagen). Equal amounts of RNA were reverse-transcribed into cDNA using iScript cDNA Synthesis Kit (Biorad). Relative gene expression was determined using gene-specific primers and SYBR Green Master Mix (Life Technologies) on a StepOne RT-PCR System (Applied Biosystems). Expression levels were normalized to *Gapdh* or *18S* using the ΔCt method or ΔΔCt relative to control groups.

### Intestinal transit assays

For all intestinal transit assays, non-fasted mice were used to preserve endogenous motility. For total GI transit, mice were gavaged with 300 ul of 6% carmine red (Sigma-Aldrich) suspended in 0.5% methylcellulose (Wako). After gavage, mice were placed in individual plastic cages and fecal pellets were monitored at 10 min intervals. Total GI transit was defined as the latency from gavage to the appearance of carmine red in fecal pellets.

For colonic transit, mice were lightly anesthetized by isoflurane and a 3 mm glass bead lubricated in 10% glycerol/water was placed onto the anal verge and inserted 2 cm into the distal colon using a marked lubricated stick. Mice were immediately placed into individual cages to awaken and the time taken to expel the glass bead was recorded.

### Intestinal permeability assay

In the morning without fasting, mice were gavaged with 200 mg/kg of fluorescein sulfonic acid (478 daltons, Thermo Fisher Scientific) dissolved in PBS as previously described (Rao et al., 2017). After 4 hr, blood was obtained by cardiac puncture, centrifuged at 2500 g for 15 min, and serum fluorescence was measured in a Synergy microplate reader (BioTek) at ex/em 490/520 nm.

### Treatments to accelerate intestinal transit

For Miralax experiments, mice had unlimited access to 10% PEG3350 (Miralax) dissolved in their drinking water continuously for 5 days prior to infection, or for 5 days during infection. Control mice received regular drinking water.

For carbachol experiments, mice were gavaged with 3.0 mg/kg of carbamoylcholine chloride (Sigma-Aldrich) dissolved in 200 ul water, followed immediately with carmine red solution to measure changes to total GI transit. During infection experiments, mice were gavaged with carbamoylcholine chloride starting 30 min prior to oral infection and continued once daily thereafter.

### 16S rRNA gene sequencing and data analysis

To analyze bacterial community structure, genomic DNA was extracted from luminal contents, mucosal scrapings, or feces. To obtain luminal contents, 5 cm of proximal duodenums, 5 cm of distal ileums, or entire colons were flushed with PBS (0.5ml PBS/cm segment) and contents were frozen at −20°C until use. To obtain mucosal scrapings, flushed tissue segments were opened longitudinally, and the mucus and associated bacteria were scraped and frozen at −20°C in 0.5 ml PBS. Freshly collected fecal pellets were frozen at −20°C until use. Upon thawing, genomic DNA was extracted using QIAamp DNA Stool Mini Kit (Qiagen). Samples were normalized to the same concentration (8 ng/ul), amplified (40 ng per reaction, 35 cycles), and barcoded in triplicate PCR reactions using 5Prime HotMaster Mix (QuantaBio). The published 515F/806R Golay-barcoded primer pair with Illumina adapters were used to amplify the V4 region of 16S rRNA gene (Caporaso et al., 2012). Barcoded triplicate reactions were combined and a 300-350 bp amplicon for each sample was confirmed by running on a DNA agarose gel. 300 ng of each sample was pooled, purified using QIAquick DNA purification column, and sequenced using 250 bp paired-end sequencing on an Illumina MiSeq (Ilumina). A mean sequence depth of 10,635 counts/sample was obtained. Paired-reads were merged, quality trimmed, and clustered into operational taxonomic units (OTUs) at 97% sequence similarity. Taxonomy was assigned using the RDP database version 11.5 (Cole et al., 2014). To estimate within-sample richness, alpha-diversity was determined using Chao1 estimates. To evaluate differences in diversity across samples, beta-diversity was determined using unweighted UniFrac distances. The relatedness of community diversity between pairs of samples was represented through principal coordinates analysis using QIIME software package (Kuczynski et al., 2011). Differences in relative abundance at different taxonomic levels were determined using ANOVA with a false discovery rate of 5% or Mann-Whitney tests.

### Fecal microbiota transplantation

For microbiota transplantation by oral gavage, fecal pellets were collected from SFB-monocolonized germ-free mice (provided by J. Huh) and frozen at −80°C until use. Alternatively, freshly collected feces were homogenized as described below and used immediately for gavage. Fresh or frozen feces were homogenized in water and passed through a 70 um strainer to remove solid debris. Mice were gavaged with 200-300 ul of cleared fecal supernatants, which corresponded to 1-2 fecal pellets per recipient mouse, twice in one day separated by 8 hr. Mice were rested for 5-10 days to allow transplanted fecal microbiota to colonize before being used for infection experiments (Farkas et al., 2015). In some experiments, mice were gavaged with their own feces as a control for fecal transplantation. In other cases, mice were gavaged with water as control since fecal transplantation could perturb the native luminal microbiota.

### Quantification of SFB

To quantify relative SFB levels, genomic DNA was extracted from ileum mucosal scrapings or feces following an established protocol (Farkas et al., 2015) with slight modifications. To obtain mucosal scrapings, the distal 5 cm of the ileum was flushed with PBS to clear luminal contents, opened longitudinally, and the mucus and associated bacteria were scraped and frozen in 0.5 ml DNA extraction buffer (140mM Tris-HCl/140mM NaCl/14mM EDTA/5.7% SDS/water) at −20°C until use. For feces, 1-2 freshly collected fecal pellets were frozen at −20°C until use. Upon thawing, mucosal scrapings and feces were homogenized in a 2 ml eppendorf containing 1 ml of DNA extraction buffer and a single 5 mm steel bead in a TissueLyzer (Qiagen) for 3 min at 30 Hz. Samples were centrifuged at 5,000 g for 5 min at 4°C. 500 ul of supernatant was transferred to a new 2 ml eppendorf tube containing 200 ul of 0.5 mm acid-washed glass beads (Sigma-Aldrich) and 500 ul of phenol:chloroform:isoamyl alcohol. Tubes were again homogenized in the TissueLyzer for 3 min at 30 Hz, then centrifuged at 16,000 g for 10 min at 4°C. 250 ul of the aqueous phase was transferred to a new eppendorf tube with 25 ul of 5M sodium acetate and 275 ul of isopropanol. Tubes were vortexed and centrifuged at 16,000 g for 10 min at 4°C to precipitate DNA. Supernatant was poured off and the DNA pellet was washed with 500 ul of 70% ethanol to remove residual salt. Tubes were centrifuged at 9,000 g for 5 min at 4°C to re-pellet DNA. Following removal of supernatant, the DNA pellet was air-dried and resuspended in 200 ul of TE buffer (10mM Tris-HCl/1mM EDTA/water). qPCR was conducted on samples to determine the amount of SFB and total Eubacteria using SYBR Green reagent and bacteria-specific primers on a StepOne RT-PCR System (Applied Biosystems).

### Microfold cell depletion

For microfold cell depletion, we followed an established protocol (Knoop et al., 2009). Mice were injected i.p. with 250 ug of anti-RANKL (clone IL22/5, BioXcell) diluted in saline every 2 days. Control mice received 250 ug of IgG2a isotype control (clone 2A3, BioXcell). Depletion of microfold cells was confirmed by immunostaining of ileal PP FAE.

### Preparation of ileum and PP epithelial cells and intraepithelial leukocytes

Ileums were flushed with ice-cold 5% FBS/PBS and the attached mesentery was removed. PP were dissected and processed separately. PPs and ileums devoid of PPs were shaken at 250 r.p.m. for 20 min at 37°C in 2mM EDTA/1mM DTT/5% FBS/RPMI to isolate epithelial cells and intraepithelial leukocytes. Tissues were vortexed and passed through a 70 um strainer to collect single cell suspensions of epithelial cells and intraepithelial leukocytes. The isolation step was repeated, and pooled cells were washed in 5% FBS/RPMI

### Preparation of ileum lamina propria leukocytes, PP immune, and PP stromal cells

After removal of epithelial cells, ileum lamina propria and PP tissues were rinsed in 5% FBS/RPMI. Ileum lamina propria was minced before digestion. Minced ileum tissues or whole PPs were digested by shaking at 250 r.p.m. for 45-60 min at 37°C in 1 mg/ml collagenase/2 mg/ml Dispase/100 U/ml DNase I/5% FBS/RPMI. Every 15 min, tissues were vortexed and the supernatant was collected by passing through a 100 um strainer. Tissues were re-incubated with fresh pre-warmed digestion media. Pooled cells were passed through a 70 um strainer to collect single cell suspensions and washed in 5% FBS/RPMI. In some cases, leukocytes were enriched using a Percoll gradient and collected at the 44:67% interphase.

### Preparation of ileum longitudinal muscle/myenteric plexus and muscularis macrophages

Ileums were flushed with ice-cold 5% FBS/PBS and the attached mesentery and PPs were removed. The longitudinal muscle layer and associated myenteric plexus were carefully separated from the mucosa/submucosa. For qRT-PCR experiments, longitudinal muscle/myenteric plexus preparations were homogenized in a 2 ml eppendorf containing 1 ml Trizol and a single 5 mm steel bead in a TissueLyzer (Qiagen) for 3 min at 30 Hz. Samples were spun down and supernatant was removed to a new eppendorf, then frozen at −80°C until RNA extraction. For flow cytometry experiments, longitudinal muscle tissues were washed, minced, and digested as described in *Preparation of ileum lamina propria and PP leukocytes*.

### Flow cytometry

For surface staining, single cell suspensions were stained for 20 min on ice with surface antibodies at 1:100 in Flow Buffer (2% BSA/0.1% sodium azide/1mM EDTA/HBSS). For Th17 cell analysis, surface antibodies included CD45-APC-Cy7 (30-F11), CD4-BV421 (RM4-5), TCRβ-FITC (H57-597) from Biolegend, and intracellular antibodies included IL-17A-PE (TC11-18H10.1) and IFNγ-APC (XMG1.2) from Biolegend. For innate lymphoid cell analysis, lineage cocktail included surface antibodies against CD3 (145-2C11), TCRδ (GL3), TCRβ (H57-597), CD11b (M1/70), CD11c (N418), CD19 (6D5), and Gr1 (RB6-8C5) in FITC channel from Biolegend, as well as CD90.1/Thy1-PerCP (OX-7), NKp46-BV421 (29A1.4), CD4-PE-Cy7 (RM4-5) from Biolegend, and RORγt-APC (BD2) from eBioscience. For muscularis macrophage analysis, surface antibodies included CD45-APC-Cy7 (30-F11), CD11b-BV605 (M1/70), CD103-PE (2E7), F480-FITC (BM8), CD64-BV421 (X54-5/7.1), and CX3CR1-PE-Cy7 (SA011F11) from Biolegend. For ileum and PP epithelial cell and intraepithelial leukocyte analysis, surface antibodies included Epcam-APC (G8.8), CD45-APC-Cy7 (30-F11), CD4-BV421 (RM4-5), CD8α-PerCP or PE-Cy-7 (53-6.7), CD8β-PE (YTS156.7.7), TCRδ-FITC or BV421 (GL3), TCRβ-FITC (H57-597), Vγ1-APC (2.11), Vδ4-PE (GL2) from Biolegend, Vγ2-PE-Cy7 (UC3-10A6) from eBioscience, UEA-1-FITC (FL-1061) from Vector Laboratories, NKM-PE (16-2-4) from Miltenyi Biotec, Vγ7-FITC (kindly provided by P. Pereira, Institut Pasteur, France), and intracellular antibody against Ki67-PE-Cy7 (SolA15) from eBioscience. For PP stromal cell analysis, surface antibodies included Epcam-APC (G8.8), CD45-APC-Cy7 (30-F11), PDPN-PE-Cy7 (8.1.1), and CD31-BV421 (390) from Biolegend. For PP immune cell analysis, surface antibodies included CD11c-PE (N418), MHC-II-PacBlue (M5/114.15.2), CD11b-BV605 (M1/70), F480-FITC (BM8), CX3CR1-PE-Cy7 (SA011F11), B220-APC (RA3-6B2), CD3-PerCP (145-2C11), CD4-BV421 (RM4-5), GL7-A488 (GL7) from Biolegend, and IgA-PE (mA-6E1) from eBioscience. Non-specific binding was blocked with mouse FcR Blocking Reagent (Miltenyi Biotec) for 20 min on ice. Dead cells were excluded using Zombie Aqua Fixable Dye (Biolegend) or Fixable Viability Dye eFlour (eBioscience). For intracellular staining of nuclear markers (RORγt and Ki67 from eBioscience), cells were stained for surface markers described above, fixed in Foxp3 Transcription Factor kit (eBioscience) for 30 min on ice, followed by incubation with intracellular antibodies at 1:100 in Permeabilization Buffer for 30 min. For intracellular staining of cytokines (IL-17A and IFNγ from Biolegend), cells were cultured for 4 hr at 37°C in 50ng/ml PMA, 1ug/ml ionomycin, and GolgiStop (BD Biosciences), stained for surface markers, fixed with Cytoperm/Cytofix Buffer (BD Biosciences) for 30 min, followed by incubation with intracellular antibodies at 1:100 in Permeabilization Buffer for 30 min. Flow cytometry was performed on a BD LSR-II flow cytometer, and data was analyzed using FlowJo software (Treestar).

### RNA-Seq analysis of ileum epithelial cells

Ileum epithelial cells were isolated as described in *Preparation of ileum and PP epithelial cells and intraepithelial leukocytes*. Cells were transferred to eppendorfs, washed twice, resuspended in 1 ml Trizol, and frozen at −80°C until RNA extraction. Total RNA was extracted by chloroform as described in *Quantitative RT-PCR* and using a PureLink RNA Mini kit (Life Technologies). RNA-Seq library preparation and sequencing by an Illumina NextSeq was conducted by the HMS Biopolymers Genomic Facility.

### Fecal IgA ELISA

Fecal pellets were collected into a 2 ml eppendorf, diluted 1:10 w/v with 1mM EDTA/PBS, homogenized with a single 5 mm steel bead in a TissueLyzer (Qiagen) at 30 Hz for 3 min. Homogenates were centrifuged twice at 12,000 g for 10 min at 4°C and supernatants collected for ELISA. NUNC Maxisorp 96-well plates (Electron Microscopy Sciences) were coated with goat anti-mouse IgA-Biotin (BioLegend) and washed with 0.05% Tween/PBS. Plates were blocked with 5% milk powder/PBS and washed. Experimental samples and standards of mouse IgA were added for 1 hr at RT. All samples were run in duplicate and were serially diluted. Plates were washed and incubated with goat IgG anti-mouse IgA-HRP (BioLegend). Plates were washed and developed in the dark using TMB substrate, then 0.18M H2SO4 was used to stop the reaction. Plates were read using a Synergy microplate reader (BioTek) at 450 nm. Fecal IgA values were quantified by comparison to standards fit to a linear regression model.

### Calcium imaging of DRG neuronal cultures

Mice were euthanized by CO_2_ inhalation, and DRGs were dissected and dissociated as previously described (Pinho-Ribeiro et al., 2018). DRG neurons were plated on laminin pre-coated 35 mm culture dishes (2,000 cells per dish) in 50 ng/ml nerve growth factor/neurobasal media (Thermo Fisher), and used for calcium imaging 12-24 hr after plating. For calcium imaging, cells were loaded with 5 mM Fura-2-AM/neurobasal media (Thermo Fisher) at 37°C for 30 min, washed twice, and imaged in 2 mL Krebs-Ringer solution (Boston Bioproducts). STm was cultured overnight as described in *Bacterial strains and culture conditions*, washed twice, resuspended in Krebs-Ringer solution (vehicle), and bath-applied at 1 × 10^9^ CFU/ml. After 200 ul of STm or vehicle stimulation, 1 uM capsaicin (Tocris) and 40 mM KCl (Sigma) was applied sequentially to identify TRPV1 and live neurons respectively. Images were acquired with alternating 340/380 nm excitation wavelengths, and fluorescence emission was captured using a Nikon Eclipse Ti inverted microscope and Zyla sCMOS camera. Ratiometric analysis of 340/380 signal intensities were processed, background corrected, and analyzed with NIS-elements software (Nikon) by drawing regions of interest (ROI) around individual cells. The percentage of STm-responsive cells with an increase in 340/380 ratio greater than 10% from 3 separate fields per condition was quantified.

### CGRP release of DRG neuronal cultures

For CGRP release, DRG neurons were dissected and cultured in 96-wells (5,000 cells per well) as previously described (Pinho-Ribeiro et al., 2018), and used in CGRP release assays 1 week after plating. One group of neurons was treated with 25 pg of Botulinum neurotoxin A (BoNT/A) for 24 hr prior to neuronal stimulation. The neuronal culture medium was removed from all wells, and 200 ul of fresh neurobasal media was added to the wells. DRG neurons were stimulated with 50 ul of live, heat-killed, filtered supernatant from STm cultures, media control (neurobasal media), or 1uM capsaicin. Live STm and filtered supernatants were prepared on the day of the experiment. STm were heat-killed by exposure to 90°C heat for 10 min. DRG neurons were stimulated for 1 hr at 37°C with 5% CO_2_, then 50 ul of supernatant were collected to determine CGRP concentration using an Enzyme Linked Immunosorbent kit (Cayman Chemical) according to the manufacturer’s instructions.

### Epithelial organoid cultures

Murine small intestinal crypts were isolated and cultured as previously described (De Lau et al., 2012; Sato et al., 2009). Briefly, ileums were flushed, opened longitudinally and cut into 2-4 mm segments. Tissues were incubated in 2 mM EDTA/PBS for 30 min on ice, pipetted, then washed to release crypts into the supernatant. Supernatant was passed through a 70 um strainer and centrifuged at 300 g for 5 min. Crypts were grown in Matrigel (Corning) in 24-well plates with Advanced DMEM/F12 cell culture media (Life Technologies) supplemented with growth factors including EGF (50 ng/ml, Life Technologies), Noggin (100 ng/ml, PeproTech), and R-spondin-1 conditioned media (supplied by Harvard Digestive Diseases Center Gastrointestinal Organoid Culture Core). For stimulation of organoids, RANKL (200 ng/ml, Biolegend) was added to wells either alone or with CGRP (1, 10, 100 nM, GenScript) and incubated for 72 hr. Fresh media, RANKL, and CGRP was replaced at 48 hr. To harvest cells, organoids were placed in Cell Recovery Solution (Corning) for 30 min prior to washing and resuspending in Trizol, then stored at −80°C prior to RNA extraction.

## Analysis of published single-cell DRG and enteric neuron expression data

Gene expression patterns of single cell RNA-seq data published by Zeisel et al. (Cell 2018) of the mouse nervous system (posted at http://mousebrain.org/genesearch.html) were analyzed for cellular subsets isolated from the dorsal root ganglia and from the enteric nervous system. DRG neuron clusters were designated by Zeisel et al. as PSPEP1-8, PSNF1-3, PSNP1-6 and enteric neuron subsets as ENT1-9. Average transcript levels of *Scn10a, Trpv1, Ntrk1, Calca, Calcb, Calb2, Hand2*, and *Phox2b* across these neuronal datasets were hierarchically clustered by Pearson correlation, average linkage, and plotted as a relative heat-map using Morpheus (Broad Institute, https://software.broadinstitute.org/morpheus/).

### Statistical analysis

Sample sizes for all experiments were chosen according to standard practice in the field. Statistical parameters, including numbers, distribution, and deviation, and statistical tests are reported in the figures and corresponding legends. Most data are represented as mean ± standard error (SEM) unless stated otherwise. Statistical analyses were performed in Microsoft Excel and GraphPad Prism.

---

**Figure S1. *Nav1.8-Cre/DTA* and RTX-treatment ablation strategies target extrinsic nociceptors and mediate host protection against *Salmonella* infection**

**(A)** Heat-map showing relative expression of distinct neural transcripts (*Scn10a, Trpv1, Ntrk1, Calca, Calcb, Calb2, Hand2, Phox2b*) from clustered populations of DRG and enteric neurons from single-cell RNA-Seq data of the mouse nervous system (Zeisel et al. 2018)

**(B, E, F)** *Nav1.8-Cre/DTA* mice (B), *Trpv1-DTR* mice (E), and RTX-treated mice (F) were orally infected with STm and sacrificed at 1 or 5 dpi to examine STm tissue burden in the cecum, mesenteric lymph nodes and Liver. See Figure 1.

**(C-D)** Quantification of TRPV1+, CGRP+, and NF200+ neurons out of total Tuj1+ neurons from lumbar DRGs and vagal ganglia of *Nav1.8-Cre/DTA* mice and control littermates. Data are pooled from Cre- (n=3) and Cre+ (n=3) mice with 2-3 fields per mouse. Scale bar, 50um.

**(G-H)** Quantification of TRPV1+, CGRP+, and NF200+ neurons out of total Tuj1+ neurons from lumbar DRGs and vagal ganglia of subcutaneous RTX-treated and vehicle-treated mice. Data are pooled from vehicle (n=4) and RTX (n=4) treated mice with 2-6 fields per mouse. Scale bar, 50um.

Statistical analysis: **(B, E, F)** Each circle represents an individual animal. Mann-Whitney tests for CFU analysis. **(C-D, G-H)** Each circle represents an individual image. Unpaired t-tests for DRG and vagal ganglia immunostaining analysis. *p<0.05, **p<0.01, ***p<0.001, ****p<0.0001. Error bars are mean ± SEM.

**Figure S2. *Nav1.8-Cre/DTA* and RTX-treatment strategies do not impact gut-intrinsic enteric neurons within the myenteric plexus**

**(A-B)** qRT-PCR of *Hand2* transcripts from DRG, vagal ganglia, ileum, and colon tissues of *Nav1.8-Cre/DTA* mice and control littermates (n=6 mice/group), or RTX-treated and vehicle-treated mice (n=5 mice/group).

**(C-E)** Quantification of HuD+, calretinin+, and neuronal nitric oxide synthase (nNOS+) neurons from myenteric plexus of *Nav1.8-Cre/DTA* mice and control littermates (n=6 mice/group). Scale bar, 80um.

(**F-H)** Quantification of HuD+, calretinin+, and nNOS+ neurons from myenteric plexus of RTX-treated mice and vehicle-treated mice (n=6 mice/group). Scale bar, 80um.

**(I)** Subcutaneous RTX-treated mice were infected intraperitoneally with 1.8 × 10^2^ CFU STm, and sacrificed at 4 dpi to examine STm tissue burden and weight loss. Data are from vehicle (n=8) and RTX (n=8) treated mice.

Statistical analysis: **(A-B)** Two–way ANOVA and Sidak test for qRT-PCR. **(E, H)** Unpaired t-tests for immunostaining of myenteric plexus. **(I)** Mann-Whitney tests for CFU analysis. Unpaired t-tests for weight loss analysis. Error bars are mean ± SEM.

**Figure S3. Nociceptor neurons do not regulate intestinal epithelial barrier or intraepithelial lymphocyte populations**

**(A)** Intestinal permeability was assessed 4 hours after gavage of fluorescent dye in *Nav1.8-Cre/DTA* mice and control littermates. Data are from uninfected mice (n=6 Cre- and 7 Cre+) and STm 2dpi mice (n=8 Cre- and 9 Cre+).

**(B)** Proportions of proliferative Ki67+Epcam+ cells in ileum epithelial cells from *Nav1.8-Cre/DTA* mice and control littermates. Data are from uninfected mice (n=6 Cre- and 6 Cre+) and STm 5dpi mice (n=6 Cre- and 6 Cre+).

**(C)** RNA-Seq analysis of ileum intestinal epithelial cells (IEC) of uninfected *Nav1.8-Cre/DTA* mice and control littermates (n=4 Cre- and 4 Cre+). Data are plotted as a volcano plot showing differentially expressed genes (>2 fold-change, p<0.05) of *Nav1.8-Cre/DTA* vs control ileal IEC transcriptomes.

**(D)** Expression of tight junction genes in ileal IEC from uninfected *Nav1.8-Cre/DTA* mice and control littermates (n=4 mice/group).

**(E)** Expression of anti-microbial genes in ileal IEC from uninfected *Nav1.8-Cre/DTA* mice and control littermates (n=4 mice/group).

**(F)** Proportions of TCRαβ and TCRγδ intraepithelial leukocytes in ileums of uninfected *Nav1.8-Cre/DTA* mice and control littermates. Data are from Cre- (n=5) and Cre+ (n=7) mice.

**(G)** Subsets of TCRγδ cells were analyzed by flow cytometry in *Nav1.8-Cre/DTA* mice and control littermates. Data are from Cre- (n=5) and Cre+ (n=6) mice.

Statistical analysis: **(A)** One-way ANOVA and Tukey post-test for intestinal permeability. **(B)** One-way ANOVA and Tukey post-test for IEC proliferation. **(D-E)** Two-way ANOVA and Tukey post-test for TJ and AMP genes. **(F-G)** Two-way ANOVA and Sidak post-test for TCRγδ IELs. Error bars are mean ± SEM.

**Figure S4. Nociceptor neurons do not regulate major immune populations in the ileum lamina propria.**

**(A)** Five days after self-feces or SFB feces transplantation, *Nav1.8-Cre/DTA* mice were orally infected with 0.34-0.91 × 10^7^ CFU STm and sacrificed at 5 dpi to analyze STm tissue burden. Data are pooled from 2 experiments with *Nav1.8-Cre*^+^/*DTA* mice gavaged with self-feces (n=13), and *Nav1.8-Cre*^+^/*DTA* mice gavaged with SFB feces (n=12).

**(B)** qPCR of SFB levels in ileum mucosa of *Nav1.8-Cre/DTA* mice gavaged with their own feces (n=4) or SFB feces (n=6) at STM 5dpi.

**(C)** Proportions of IL-17A+ and IFNγ+ cells out of CD4+TCRβ+ T cells in ileum lamina propria of *Nav1.8-Cre/DTA* mice. For Th17 analysis, data are pooled from 2 experiments with uninfected mice (n=5 Cre- and 6 Cre+) and STM 2dpi mice (n=7 Cre- and 9 Cre+). For Th1 analysis, data are pooled from 3 experiments with uninfected mice (n=8 Cre- and 9 Cre+) and STm 2dpi mice (n=7 Cre- and 9 Cre+).

**(D)** Proportions of UEA-1+ cells out of Epcam+ cells in ileum epithelial cells from *Nav1.8-Cre/DTA* mice and control littermates. Data are pooled from 2 experiments with uninfected mice (n=6 Cre- and 6 Cre+) and STm 5dpi mice (n=6 Cre- and 6 Cre+).

**(E)** Proportions of Nkp46+, CD4+RORγt+ and CD4-RORγt+ cells out of Lin-Thy1+ innate lymphoid cells in ileum lamina propria from uninfected *Nav1.8-Cre/DTA* mice and control littermates. Data are from Cre- (n=4) and Cre+ (n=3) mice.

**(F)** Proportions of CD11b+F480+CD64+CX3CR1+ macrophages in ileum muscularis from *Nav1.8-Cre/DTA* mice and control littermates. Data are from 2 experiments with uninfected mice (n=5 Cre- and 7 Cre+) and STm 5dpi mice (n=4 Cre- and 7 Cre+).

Statistical analysis: **(A)** Mann-Whitney tests for CFU analysis. **(B)** Mann-Whitney tests for SFB qPCR. **(C,D,F)** One-way ANOVA and Sidak post-tests for T cells, fucosylated epithelial cells, and muscularis macrophages. **(E)** Unpaired t-tests for ILC analysis. *p<0.05, **p<0.01. Error bars are mean ± SEM.

**Figure S5. Nociceptor regulation of gastrointestinal transit and impact on host defense**

**(A)** Total gastrointestinal (GI) transit, as measured by time until carmine red dye appears in feces, in nociceptor-ablated mice. Data are pooled from 2 experiments from *Nav1.8-Cre/DTA* mice and control littermates (n=9 Cre- and n=13 Cre+), 3 experiments for subcutaneous vehicle and RTX-treated mice (n=22 vehicle and n=23 RTX), and 4 experiments for intrathecal vehicle and RTX-treated mice (n=19 vehicle and n=19 RTX).

**(B)** Colon transit, as measured by time taken for bead to be expelled after being pushed 2 cm into the colon, in nociceptor-ablated mice. Data are from *Nav1.8-Cre/DTA* mice and control littermates (n=13 Cre- and 12 Cre+), subcutaneous vehicle and RTX-treated mice (n=9 vehicle and 10 RTX), and intrathecal vehicle and RTX-treated mice (n=8 vehicle and 8 RTX).

**(C)** Total GI transit was measured in subcutaneously injected vehicle and RTX-treated mice at baseline and after administration of 10% Miralax in drinking water. Data are from water-treated mice (n=6 vehicle and 6 RTX) and Miralax-treated mice (n=6 vehicle and 6 RTX).

**(D)** Subcutaneously injected vehicle and RTX-treated mice were administered with 10% Miralax in their drinking water or regular drinking water during infection. Mice were orally infected with 0.93 × 10^7^ CFU STm and sacrificed at 5 dpi to analyze STm load in ileum and spleen. Data are from water-treated mice (n=5 vehicle and 5 RTX) and Miralax-treated mice (n=6 vehicle and 5 RTX).

**(E)** Subcutaneously injected vehicle and RTX-treated mice were administered with 10% Miralax in their drinking water for 5 days prior to STm infection. On the day of infection, mice were switched to regular water, orally infected with 0.93 × 10^7^ CFU STm, and sacrificed at 5 dpi to analyze STm load in ileum and spleen. Data are from water-treated mice (n=5 vehicle and 5 RTX) and Miralax-treated mice (n=5 vehicle and 5 RTX).

**(F)** Total GI transit in wildtype mice gavaged with saline or carbachol (3mg/kg). Data are from saline (n=4) and carbachol (n=4) treated mice.

**(G)** Wildtype mice gavaged daily with saline or carbachol (3mg/kg) were infected with 1.6 × 10^7^ CFU STm, and sacrificed at 5 dpi to analyze STm load in Peyer’s patch and spleen. Data are from saline (n=4) and carbachol (n=4) treated mice.

Statistical analysis: **(A-B, F)** Unpaired t-tests for total GI transit and colon transit. **(C)** Two-way ANOVA for total GI transit. **(D-E)** Non-parametric one-way ANOVA and Dunn’s post-test for CFU analysis. **(G)** Mann-Whitney tests for CFU analysis. *p<0.05, **p<0.01, ***p<0.001. Error bars are mean ± SEM.

**Figure S6. Analysis of Peyer’s Patch numbers, epithelial, stromal, and immune cell populations in nociceptor-ablated mice**

**(A)** *Nav1.8-Cre/DTA* mice and control littermates were infected with 0.8 × 10^7^ CFU of wildtype STm or 0.9 × 10^7^ CFU mutant Δ*FimH* STm and sacrificed at 1 dpi to analyze bacterial load in the cecum. See also Figure 5B.

**(B)** Number of Peyer’s patches in the duodenum, jejunum, and ileum of the small intestine of *Nav1.8-Cre/DTA* mice and control littermates (n=23 Cre- and 20 Cre+).

**(C)** Number of follicles per Peyer’s Patch from *Nav1.8-Cre/DTA* mice and control littermates (n=32 Cre- and 33 Cre+).

**(D)** Follicle area in ileal PPs of *Nav1.8-Cre/DTA* mice and control littermates (n=30 Cre- and 24 Cre+).

**(E)** Proportions of proliferative Ki67+Epcam+ epithelial cells from PP of RTX-treated mice. Data are from 2 experiments with vehicle (n=6) and RTX (n=6) treated mice.

**(F)** qRT-PCR of RANKL (*Tnfsfr11a*) and OPG (*Tnfsfr11b*) transcripts in whole ileal PP tissues of RTX-treated mice. Data are from 2 experiments with vehicle (n=6) and RTX (n=6) treated mice.

**(G)** Proportions of stromal cells in ileal PPs of RTX-treated mice. Data are from 3 experiments with vehicle (n=9) and RTX (n=7) treated mice. Fibroblast reticular cells (FRC): PDPN+CD31-, lymphatic endothelial cells (LEC): PDPN+CD31+, and blood endothelial cells (BEC): CD31+PDPN-.

**(H)** Proportions of CD11b+ dendritic cells (CD11c+MHC-2+), CD11b-dendritic cells, and F480+ macrophages (CD3-CD19-CD11c-MHC-2+) in ileal PPs of RTX-treated mice. Data are from 3 experiments with vehicle (n=11) and RTX (n=11) treated mice.

**(I)** Proportions of CD3+ T cells, CD4+ and CD4-T cells in ileal PPs of RTX-treated mice. Data are from 3 experiments with vehicle (n=11) and RTX (n=11) treated mice.

**(J)** Proportion of B220+ B cells in ileal PPs of RTX-treated mice. Data are from 2 experiments with vehicle (n=8) and RTX (n=8) treated mice.

**(K)** Proportion of IgA+ plasma B cells and GL7+ germinal center B cells in ileal PPs of RTX-treated mice. Data are from 2 experiments with vehicle (n=8) and RTX (n=8) treated mice.

Statistical analysis: **(A)** Non-parametric one-way ANOVA and Dunn’s post-tests for CFU analysis**. (B-D, F-K)** Unpaired t-tests for PP number, follicle number, follicle area, ileal PP qRT-PCR, PP stromal cells, myeloid cells, T cells, B cells, and B cell subsets. **(E)** One-way ANOVA and Sidak post-test for PP epithelial proliferation. *p<0.05, **p<0.01. Error bars are mean ± SEM.

**Figure S7. Role of CGRP in modulating M cells and *Salmonella* infection**

**(A)** Quantification of GP2 density in ileal PP FAE of RTX-treated mice at STm 5dpi.

Data are represented as GP2 density per follicle (left), or GP2 density of 3-6 follicles averaged per mouse (right). Data are from vehicle (n=4) and RTX (n=3) treated mice. Scale bar, 100um.

**(B)** Fecal IgA levels in uninfected nociceptor-ablated mice. Data are from RTX-treated mice (n=7 vehicle and 7 RTX), or *Nav1.8-Cre/DTA* mice (n=7 Cre- and 10 Cre+).

**(C)** Quantification of proportions of STm-responsive neurons out of total neurons (KCl-responsive) or TRPV1+ neurons (Capsaicin-responsive) by calcium imaging (n=5 fields/condition).

**(D)** qRT-PCR of *Calca* (n=6 mice/group) and *Calcb* transcripts (n=3 mice/group) from DRG, vagal ganglia, ileum, and colon tissues of *Nav1.8-Cre/DTA* mice and control littermates.

**(E)** Quantification of GP2 density in ileal PP FAE from uninfected *Calcb+/+* and *Calcb*-/-littermates. Data are from *Calcb*^+/+^ (n=3) and *Calcb*^−/−^ (n=5) mice. Scale bar, 100um.

**(F)** qRT-PCR of *Gp2* and *Spib* transcripts from small intestine epithelial organoids stimulated with RANKL (10nM) and CGRP (1, 10, 100nM) (n=3 samples/condition).

Statistical analysis: **(A-C,E)** Unpaired t-tests for GP2 analysis, fecal IgA and calcium imaging. **(D)** Two–way ANOVA and Sidak post-tests for qRT-PCR. **(F)** One-way ANOVA and Sidak post-tests for organoid qRT-PCR. *p<0.05, **p<0.01, ***p<0.001. Error bars are mean ± SEM.

**Summary Figure.**
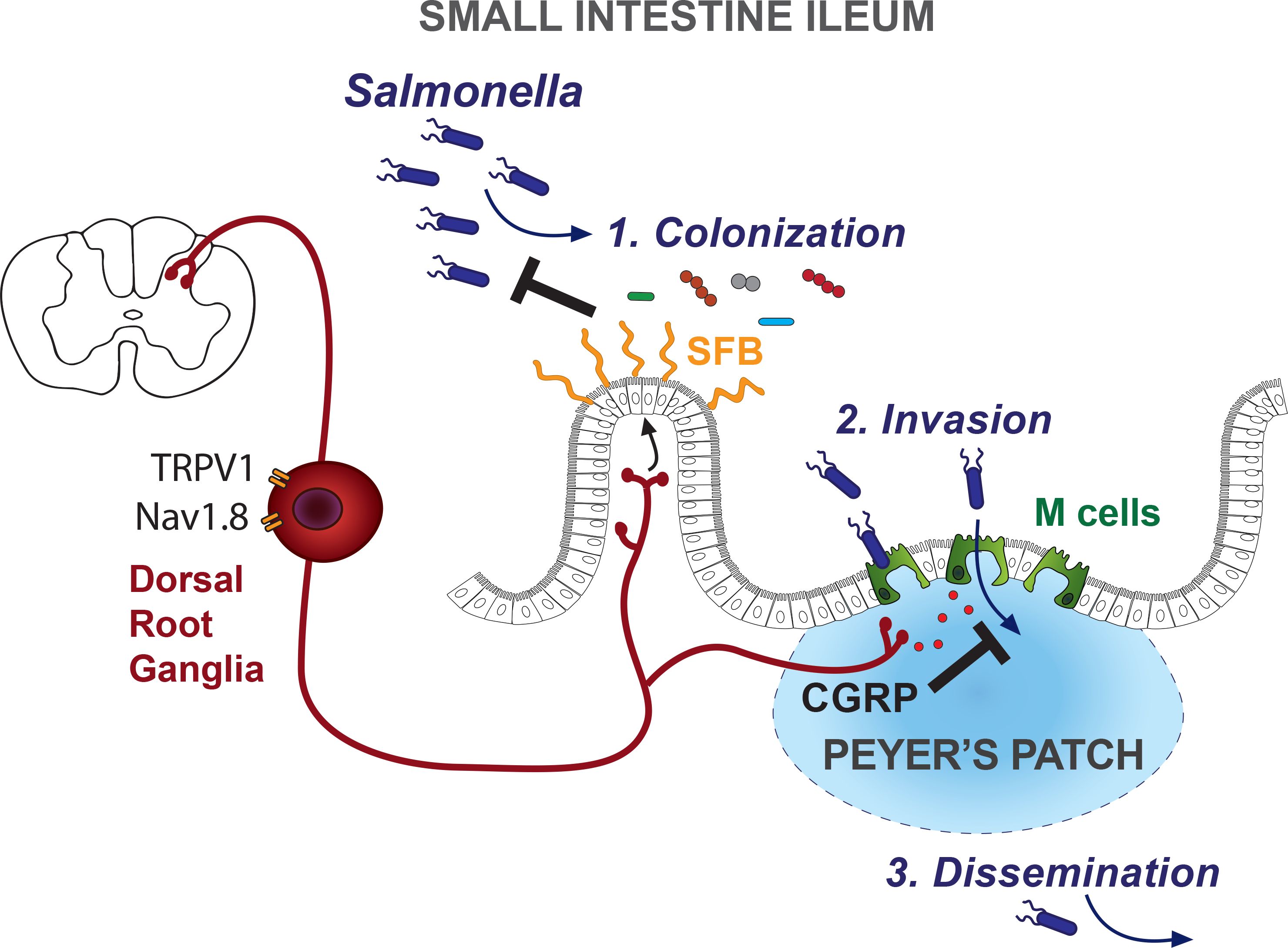
Nociceptor neurons modulate ileal microbial homeostasis and Peyer’s patch M cells to protect the host against *Salmonella* infection. Nav1.8 and TRPV1 nociceptor neurons from gut-extrinsic sensory ganglia are critical in protecting the host against *Salmonella typhimurium* (STm) infection. Nociceptors regulated the gut microbiota at homeostasis, specifically Segmented Filamentous Bacteria (SFB) levels in the ileum, which protected against STm by colonization resistance. Nociceptors also regulated the density of Microfold cells (M cells) in Peyer’s patch epithelia via CGRP to limit entry points for STm invasion into host tissues. As a result, nociceptors protected the host against subsequent STm dissemination to peripheral tissues.

## REFERENCES

Abraham, C., and Medzhitov, R. (2011). Interactions between the host innate immune system and microbes in inflammatory bowel disease. Gastroenterology 140, 1729–1737.

Abrahamsen, B., Zhao, J., Asante, C.O., Cendan, C.M., Marsh, S., Martinez-Barbera, J.P., Nassar, M.A., Dickenson, A.H., and Wood, J.N. (2008). The cell and molecular basis of mechanical, cold, and inflammatory pain. Science 321, 702–705.

Amaya, F., Decosterd, I., Samad, T.A., Plumpton, C., Tate, S., Mannion, R.J., Costigan, M., and Woolf, C.J. (2000). Diversity of expression of the sensory neuron-specific TTX-resistant voltage-gated sodium ion channels SNS and SNS2. Mol Cell Neurosci 15, 331–342.

Andrews, J.R., Qamar, F.N., Charles, R.C., and Ryan, E.T. (2018). Extensively Drug-Resistant Typhoid - Are Conjugate Vaccines Arriving Just in Time? N Engl J Med 379, 1493–1495.

Assas, B.M., Pennock, J.I., and Miyan, J.A. (2014). Calcitonin gene-related peptide is a key neurotransmitter in the neuro-immune axis. Front Neurosci 8, 23.

Atarashi, K., Tanoue, T., Ando, M., Kamada, N., Nagano, Y., Narushima, S., Suda, W., Imaoka, A., Setoyama, H., Nagamori, T., et al. (2015). Th17 Cell Induction by Adhesion of Microbes to Intestinal Epithelial Cells. Cell 163, 367–380.

Augustyniak, D., Nowak, J., and Lundy, F.T. (2012). Direct and indirect antimicrobial activities of neuropeptides and their therapeutic potential. Curr Protein Pept Sci 13, 723–738.

Baral, P., Umans, B.D., Li, L., Wallrapp, A., Bist, M., Kirschbaum, T., Wei, Y., Zhou, Y., Kuchroo, V.K., Burkett, P.R., et al. (2018). Nociceptor sensory neurons suppress neutrophil and gammadelta T cell responses in bacterial lung infections and lethal pneumonia. Nat Med 24, 417–426.

Bartho, L., Benko, R., Holzer-Petsche, U., Holzer, P., Undi, S., and Wolf, M. (2008). Role of extrinsic afferent neurons in gastrointestinal motility. Eur Rev Med Pharmacol Sci 12 Suppl 1, 21–31.

Basbaum, A.I., Bautista, D.M., Scherrer, G., and Julius, D. (2009). Cellular and molecular mechanisms of pain. Cell 139, 267–284.

Baumler, A.J., and Sperandio, V. (2016). Interactions between the microbiota and pathogenic bacteria in the gut. Nature 535, 85–93.

Behnsen, J., Perez-Lopez, A., Nuccio, S.P., and Raffatellu, M. (2015). Exploiting host immunity: the Salmonella paradigm. Trends Immunol 36, 112–120.

Benarroch, E.E. (2015). Ion channels in nociceptors: recent developments. Neurology 84, 1153–1164.

Bioley, G., Monnerat, J., Lotscher, M., Vonarburg, C., Zuercher, A., and Corthesy, B. (2017). Plasma-Derived Polyreactive Secretory-Like IgA and IgM Opsonizing Salmonella enterica Typhimurium Reduces Invasion and Gut Tissue Inflammation through Agglutination. Front Immunol 8, 1043.

Blackshaw, L.A., Brookes, S.J., Grundy, D., and Schemann, M. (2007). Sensory transmission in the gastrointestinal tract. Neurogastroenterol Motil 19, 1–19.

Blackshaw, L.A., and Gebhart, G.F. (2002). The pharmacology of gastrointestinal nociceptive pathways. Curr Opin Pharmacol 2, 642–649.

Blake, K.J., Baral, P., Voisin, T., Lubkin, A., Pinho-Ribeiro, F.A., Adams, K.L., Roberson, D.P., Ma, Y.C., Otto, M., Woolf, C.J., et al. (2018). Staphylococcus aureus produces pain through pore-forming toxins and neuronal TRPV1 that is silenced by QX-314. Nature Communications 9, 37.

Blander, J.M., Longman, R.S., Iliev, I.D., Sonnenberg, G.F., and Artis, D. (2017). Regulation of inflammation by microbiota interactions with the host. Nat Immunol 18, 851–860.

Boesmans, W., Owsianik, G., Tack, J., Voets, T., and Vanden Berghe, P. (2011). TRP channels in neurogastroenterology: opportunities for therapeutic intervention. Br J Pharmacol 162, 18–37.

Bolotin, A., de Wouters, T., Schnupf, P., Bouchier, C., Loux, V., Rhimi, M., Jamet, A., Dervyn, R., Boudebbouze, S., Blottiere, H.M., et al. (2014). Genome Sequence of “Candidatus Arthromitus” sp. Strain SFB-Mouse-NL, a Commensal Bacterium with a Key Role in Postnatal Maturation of Gut Immune Functions. Genome Announc 2.

Boyce, B.F., and Xing, L. (2007). Biology of RANK, RANKL, and osteoprotegerin. Arthritis Res Ther 9 Suppl 1, S1.

Broz, P., Ohlson, M.B., and Monack, D.M. (2012). Innate immune response to Salmonella typhimurium, a model enteric pathogen. Gut Microbes 3, 62–70.

Buffie, C.G., and Pamer, E.G. (2013). Microbiota-mediated colonization resistance against intestinal pathogens. Nat Rev Immunol 13, 790–801.

Caporaso, J.G., Lauber, C.L., Walters, W.A., Berg-Lyons, D., Huntley, J., Fierer, N., Owens, S.M., Betley, J., Fraser, L., Bauer, M., et al. (2012). Ultra-high-throughput microbial community analysis on the Illumina HiSeq and MiSeq platforms. ISME J 6, 1621–1624.

Cardoso, V., Chesne, J., Ribeiro, H., Garcia-Cassani, B., Carvalho, T., Bouchery, T., Shah, K., Barbosa-Morais, N.L., Harris, N., and Veiga-Fernandes, H. (2017). Neuronal regulation of type 2 innate lymphoid cells via neuromedin U. Nature 549, 277–281.

Carter, P.B., and Collins, F.M. (1974). The route of enteric infection in normal mice. J Exp Med 139, 1189–1203.

Cervi, A.L., Lukewich, M.K., and Lomax, A.E. (2014). Neural regulation of gastrointestinal inflammation: role of the sympathetic nervous system. Auton Neurosci 182, 83–88.

Chavan, S.S., Pavlov, V.A., and Tracey, K.J. (2017). Mechanisms and Therapeutic Relevance of Neuro-immune Communication. Immunity 46, 927–942.

Chiocchetti, R., Mazzuoli, G., Albanese, V., Mazzoni, M., Clavenzani, P., Lalatta-Costerbosa, G., Lucchi, M.L., Di Guardo, G., Marruchella, G., and Furness, J.B. (2008). Anatomical evidence for ileal Peyer’s patches innervation by enteric nervous system: a potential route for prion neuroinvasion? Cell Tissue Res 332, 185–194.

Chiu, I.M., Heesters, B.A., Ghasemlou, N., Von Hehn, C.A., Zhao, F., Tran, J., Wainger, B., Strominger, A., Muralidharan, S., Horswill, A.R., et al. (2013). Bacteria activate sensory neurons that modulate pain and inflammation. Nature 501, 52–57.

Coburn, B., Grassl, G.A., and Finlay, B.B. (2007). Salmonella, the host and disease: a brief review. Immunol Cell Biol 85, 112–118.

Cole, J.R., Wang, Q., Fish, J.A., Chai, B., McGarrell, D.M., Sun, Y., Brown, C.T., Porras-Alfaro, A., Kuske, C.R., and Tiedje, J.M. (2014). Ribosomal Database Project: data and tools for high throughput rRNA analysis. Nucleic Acids Res 42, D633–642.

D’Autreaux, F., Morikawa, Y., Cserjesi, P., and Gershon, M.D. (2007). Hand2 is necessary for terminal differentiation of enteric neurons from crest-derived precursors but not for their migration into the gut or for formation of glia. Development 134, 2237–2249.

De Lau, W., Kujala, P., Schneeberger, K., Middendorp, S., Li, V.S., Barker, N., Martens, A., Hofhuis, F., DeKoter, R.P., Peters, P.J., et al. (2012). Peyer’s patch M cells derived from Lgr5(+) stem cells require SpiB and are induced by RankL in cultured “miniguts”. Mol Cell Biol 32, 3639–3647.

De Schepper, S., Verheijden, S., Aguilera-Lizarraga, J., Viola, M.F., Boesmans, W., Stakenborg, N., Voytyuk, I., Schmidt, I., Boeckx, B., Dierckx de Casterle, I., et al. (2018). Self-Maintaining Gut Macrophages Are Essential for Intestinal Homeostasis. Cell 175, 400–415 e413.

Di Giovangiulio, M., Verheijden, S., Bosmans, G., Stakenborg, N., Boeckxstaens, G.E., and Matteoli, G. (2015). The Neuromodulation of the Intestinal Immune System and Its Relevance in Inflammatory Bowel Disease. Front Immunol 6, 590.

Donaldson, D.S., Sehgal, A., Rios, D., Williams, I.R., and Mabbott, N.A. (2016a). Increased Abundance of M Cells in the Gut Epithelium Dramatically Enhances Oral Prion Disease Susceptibility. PLoS Pathog 12, e1006075.

Donaldson, G.P., Lee, S.M., and Mazmanian, S.K. (2016b). Gut biogeography of the bacterial microbiota. Nat Rev Microbiol 14, 20–32.

Duerkop, B.A., Vaishnava, S., and Hooper, L.V. (2009). Immune responses to the microbiota at the intestinal mucosal surface. Immunity 31, 368–376.

El Karim, I.A., Linden, G.J., Orr, D.F., and Lundy, F.T. (2008). Antimicrobial activity of neuropeptides against a range of micro-organisms from skin, oral, respiratory and gastrointestinal tract sites. J Neuroimmunol 200, 11–16.

Elekes, K., Helyes, Z., Nemeth, J., Sandor, K., Pozsgai, G., Kereskai, L., Borzsei, R., Pinter, E., Szabo, A., and Szolcsanyi, J. (2007). Role of capsaicin-sensitive afferents and sensory neuropeptides in endotoxin-induced airway inflammation and consequent bronchial hyperreactivity in the mouse. Regul Pept 141, 44–54.

Evangelista, S. (2014). Capsaicin Receptor as Target of Calcitonin Gene-Related Peptide in the Gut. In Capsaicin as a Therapeutic Molecule, O.M.E. Abdel-Salam, ed. (Basel: Springer Basel), pp. 259–276.

Farkas, A.M., Panea, C., Goto, Y., Nakato, G., Galan-Diez, M., Narushima, S., Honda, K., and Ivanov, II (2015). Induction of Th17 cells by segmented filamentous bacteria in the murine intestine. J Immunol Methods 421, 104–111.

Furness, J.B., Nguyen, T.V., Nurgali, K., and Shimizu, Y. (2009). The Enteric Nervous System and Its Extrinsic Connections. In Textbook of Gastroenterology (Blackwell Publishing), pp. 15–39.

Gabanyi, I., Muller, P.A., Feighery, L., Oliveira, T.Y., Costa-Pinto, F.A., and Mucida, D. (2016). Neuro-immune Interactions Drive Tissue Programming in Intestinal Macrophages. Cell 164, 378–391.

Gaboriau-Routhiau, V., Rakotobe, S., Lecuyer, E., Mulder, I., Lan, A., Bridonneau, C., Rochet, V., Pisi, A., De Paepe, M., Brandi, G., et al. (2009). The key role of segmented filamentous bacteria in the coordinated maturation of gut helper T cell responses. Immunity 31, 677–689.

Garland, C.D., Lee, A., and Dickson, M.R. (1982). Segmented filamentous bacteria in the rodent small intestine: Their colonization of growing animals and possible role in host resistance to Salmonella. Microb Ecol 8, 181–190.

Garrett, W.S., Gordon, J.I., and Glimcher, L.H. (2010). Homeostasis and inflammation in the intestine. Cell 140, 859–870.

Gorkiewicz, G., Thallinger, G.G., Trajanoski, S., Lackner, S., Stocker, G., Hinterleitner, T., Gully, C., and Hogenauer, C. (2013). Alterations in the colonic microbiota in response to osmotic diarrhea. PLoS One 8, e55817.

Goto, Y., Obata, T., Kunisawa, J., Sato, S., Ivanov, II, Lamichhane, A., Takeyama, N., Kamioka, M., Sakamoto, M., Matsuki, T., et al. (2014). Innate lymphoid cells regulate intestinal epithelial cell glycosylation. Science 345, 1254009.

Grundy, L., Erickson, A., and Brierley, S.M. (2018). Visceral Pain. Annu Rev Physiol.

Haber, A.L., Biton, M., Rogel, N., Herbst, R.H., Shekhar, K., Smillie, C., Burgin, G., Delorey, T.M., Howitt, M.R., Katz, Y., et al. (2017). A single-cell survey of the small intestinal epithelium. Nature 551, 333–339.

Hase, K., Kawano, K., Nochi, T., Pontes, G.S., Fukuda, S., Ebisawa, M., Kadokura, K., Tobe, T., Fujimura, Y., Kawano, S., et al. (2009). Uptake through glycoprotein 2 of FimH(+) bacteria by M cells initiates mucosal immune response. Nature 462, 226–230.

Heczko, U., Abe, A., and Finlay, B.B. (2000). Segmented filamentous bacteria prevent colonization of enteropathogenic Escherichia coli O103 in rabbits. J Infect Dis 181, 1027–1033.

Hodges, K., and Gill, R. (2010). Infectious diarrhea: Cellular and molecular mechanisms. Gut Microbes 1, 4–21.

Holzer, P. (2002). Sensory neurone responses to mucosal noxae in the upper gut: relevance to mucosal integrity and gastrointestinal pain. Neurogastroenterol Motil 14, 459–475.

Hughes, D.T., and Sperandio, V. (2008). Inter-kingdom signalling: communication between bacteria and their hosts. Nat Rev Microbiol 6, 111–120.

Ichikawa, T., Ishihara, K., Kusakabe, T., Hiruma, H., Kawakami, T., and Hotta, K. (2000). CGRP modulates mucin synthesis in surface mucus cells of rat gastric oxyntic mucosa. Am J Physiol Gastrointest Liver Physiol 279, G82–89.

Ismail, A.S., Severson, K.M., Vaishnava, S., Behrendt, C.L., Yu, X., Benjamin, J.L., Ruhn, K.A., Hou, B., DeFranco, A.L., Yarovinsky, F., et al. (2011). Gammadelta intraepithelial lymphocytes are essential mediators of host-microbial homeostasis at the intestinal mucosal surface. Proc Natl Acad Sci U S A 108, 8743–8748.

Ivanov, II, Atarashi, K., Manel, N., Brodie, E.L., Shima, T., Karaoz, U., Wei, D., Goldfarb, K.C., Santee, C.A., Lynch, S.V., et al. (2009). Induction of intestinal Th17 cells by segmented filamentous bacteria. Cell 139, 485–498.

Jang, M.H., Kweon, M.N., Iwatani, K., Yamamoto, M., Terahara, K., Sasakawa, C., Suzuki, T., Nochi, T., Yokota, Y., Rennert, P.D., et al. (2004). Intestinal villous M cells: an antigen entry site in the mucosal epithelium. Proc Natl Acad Sci U S A 101, 6110–6115.

Jepson, M.A., and Clark, M.A. (2001). The role of M cells in Salmonella infection. Microbes Infect 3, 1183–1190.

Jones, B.D., Ghori, N., and Falkow, S. (1994). Salmonella typhimurium initiates murine infection by penetrating and destroying the specialized epithelial M cells of the Peyer’s patches. J Exp Med 180, 15–23.

Julius, D. (2013). TRP channels and pain. Annu Rev Cell Dev Biol 29, 355–384.

Kamada, N., Seo, S.U., Chen, G.Y., and Nunez, G. (2013). Role of the gut microbiota in immunity and inflammatory disease. Nat Rev Immunol 13, 321–335.

Kanaya, T., Hase, K., Takahashi, D., Fukuda, S., Hoshino, K., Sasaki, I., Hemmi, H., Knoop, K.A., Kumar, N., Sato, M., et al. (2012). The Ets transcription factor Spi-B is essential for the differentiation of intestinal microfold cells. Nat Immunol 13, 729–736.

Kashem, S.W., Riedl, M.S., Yao, C., Honda, C.N., Vulchanova, L., and Kaplan, D.H. (2015). Nociceptive Sensory Fibers Drive Interleukin-23 Production from CD301b+ Dermal Dendritic Cells and Drive Protective Cutaneous Immunity. Immunity 43, 515–526.

Klose, C.S.N., Mahlakoiv, T., Moeller, J.B., Rankin, L.C., Flamar, A.L., Kabata, H., Monticelli, L.A., Moriyama, S., Putzel, G.G., Rakhilin, N., et al. (2017). The neuropeptide neuromedin U stimulates innate lymphoid cells and type 2 inflammation. Nature 549, 282–286.

Knoop, K.A., Kumar, N., Butler, B.R., Sakthivel, S.K., Taylor, R.T., Nochi, T., Akiba, H., Yagita, H., Kiyono, H., and Williams, I.R. (2009). RANKL Is Necessary and Sufficient to Initiate Development of Antigen-Sampling M Cells in the Intestinal Epithelium. The Journal of Immunology 183, 5738–5747.

Knoop, K.A., and Newberry, R.D. (2018). Goblet cells: multifaceted players in immunity at mucosal surfaces. Mucosal Immunology 11, 1551–1557.

Kobayashi, A., Donaldson, D.S., Erridge, C., Kanaya, T., Williams, I.R., Ohno, H., Mahajan, A., and Mabbott, N.A. (2013). The functional maturation of M cells is dramatically reduced in the Peyer’s patches of aged mice. Mucosal Immunol 6, 1027–1037.

Kuczynski, J., Stombaugh, J., Walters, W.A., Gonzalez, A., Caporaso, J.G., and Knight, R. (2011). Using QIIME to analyze 16S rRNA gene sequences from microbial communities. Curr Protoc Bioinformatics Chapter 10, Unit 10 17.

Kulkarni, D.H., McDonald, K.G., Knoop, K.A., Gustafsson, J.K., Kozlowski, K.M., Hunstad, D.A., Miller, M.J., and Newberry, R.D. (2018). Goblet cell associated antigen passages are inhibited during Salmonella typhimurium infection to prevent pathogen dissemination and limit responses to dietary antigens. Mucosal Immunol 11, 1103–1113.

Lai, J., Porreca, F., Hunter, J.C., and Gold, M.S. (2004). Voltage-gated sodium channels and hyperalgesia. Annu Rev Pharmacol Toxicol 44, 371–397.

Littman, D.R., and Pamer, E.G. (2011). Role of the commensal microbiota in normal and pathogenic host immune responses. Cell Host Microbe 10, 311–323.

Liu, T., Xu, Z.-Z., Park, C.-K., Berta, T., and Ji, R.-R. (2010). Toll-like receptor 7 mediates pruritus. Nature Neuroscience 13, 1460.

Mabbott, N.A., Donaldson, D.S., Ohno, H., Williams, I.R., and Mahajan, A. (2013). Microfold (M) cells: important immunosurveillance posts in the intestinal epithelium. Mucosal Immunol 6, 666–677.

Madisen, L., Zwingman, T.A., Sunkin, S.M., Oh, S.W., Zariwala, H.A., Gu, H., Ng, L.L., Palmiter, R.D., Hawrylycz, M.J., Jones, A.R., et al. (2010). A robust and high-throughput Cre reporting and characterization system for the whole mouse brain. Nat Neurosci 13, 133–140.

Madison, B.B., Dunbar, L., Qiao, X.T., Braunstein, K., Braunstein, E., and Gumucio, D.L. (2002). Cis elements of the villin gene control expression in restricted domains of the vertical (crypt) and horizontal (duodenum, cecum) axes of the intestine. J Biol Chem 277, 33275–33283.

Majowicz, S.E., Musto, J., Scallan, E., Angulo, F.J., Kirk, M., O’Brien, S.J., Jones, T.F., Fazil, A., Hoekstra, R.M., and International Collaboration on Enteric Disease ’Burden of Illness, S. (2010). The global burden of nontyphoidal Salmonella gastroenteritis. Clin Infect Dis 50, 882–889.

Margolis, K.G., Gershon, M.D., and Bogunovic, M. (2016). Cellular Organization of Neuroimmune Interactions in the Gastrointestinal Tract. Trends Immunol 37, 487–501.

Maruyama, K., Takayama, Y., Kondo, T., Ishibashi, K.I., Sahoo, B.R., Kanemaru, H., Kumagai, Y., Martino, M.M., Tanaka, H., Ohno, N., et al. (2017). Nociceptors Boost the Resolution of Fungal Osteoinflammation via the TRP Channel-CGRP-Jdp2 Axis. Cell Rep 19, 2730–2742.

McGhie, E.J., Brawn, L.C., Hume, P.J., Humphreys, D., and Koronakis, V. (2009). Salmonella takes control: effector-driven manipulation of the host. Curr Opin Microbiol 12, 117–124.

Meseguer, V., Alpizar, Y.A., Luis, E., Tajada, S., Denlinger, B., Fajardo, O., Manenschijn, J.A., Fernandez-Pena, C., Talavera, A., Kichko, T., et al. (2014). TRPA1 channels mediate acute neurogenic inflammation and pain produced by bacterial endotoxins. Nat Commun 5, 3125.

Miller, H., Zhang, J., Kuolee, R., Patel, G.B., and Chen, W. (2007). Intestinal M cells: the fallible sentinels? World J Gastroenterol 13, 1477–1486.

Mishra, S.K., and Hoon, M.A. (2010). Ablation of TrpV1 neurons reveals their selective role in thermal pain sensation. Mol Cell Neurosci 43, 157–163.

Mishra, S.K., Tisel, S.M., Orestes, P., Bhangoo, S.K., and Hoon, M.A. (2011). TRPV1-lineage neurons are required for thermal sensation. EMBO J 30, 582–593.

Monack, D.M., Hersh, D., Ghori, N., Bouley, D., Zychlinsky, A., and Falkow, S. (2000). Salmonella exploits caspase-1 to colonize Peyer’s patches in a murine typhoid model. J Exp Med 192, 249–258.

Mulderry, P.K., Ghatei, M.A., Spokes, R.A., Jones, P.M., Pierson, A.M., Hamid, Q.A., Kanse, S., Amara, S.G., Burrin, J.M., Legon, S., et al. (1988). Differential expression of alpha-CGRP and beta-CGRP by primary sensory neurons and enteric autonomic neurons of the rat. Neuroscience 25, 195–205.

Muller, P.A., Koscso, B., Rajani, G.M., Stevanovic, K., Berres, M.L., Hashimoto, D., Mortha, A., Leboeuf, M., Li, X.M., Mucida, D., et al. (2014). Crosstalk between Muscularis Macrophages and Enteric Neurons Regulates Gastrointestinal Motility. Cell 158, 1210.

Nagashima, K., Sawa, S., Nitta, T., Tsutsumi, M., Okamura, T., Penninger, J.M., Nakashima, T., and Takayanagi, H. (2017). Identification of subepithelial mesenchymal cells that induce IgA and diversify gut microbiota. Nat Immunol 18, 675–682.

Oh-hashi, Y., Shindo, T., Kurihara, Y., Imai, T., Wang, Y., Morita, H., Imai, Y., Kayaba, Y., Nishimatsu, H., Suematsu, Y., et al. (2001). Elevated sympathetic nervous activity in mice deficient in alphaCGRP. Circ Res 89, 983–990.

Ohno, H. (2016). Intestinal M cells. J Biochem 159, 151–160.

Pinho-Ribeiro, F.A., Baddal, B., Haarsma, R., O’Seaghdha, M., Yang, N.J., Blake, K.J., Portley, M., Verri, W.A., Dale, J.B., Wessels, M.R., et al. (2018). Blocking Neuronal Signaling to Immune Cells Treats Streptococcal Invasive Infection. Cell 173, 1083–1097 e1022.

Plaisancie, P., Barcelo, A., Moro, F., Claustre, J., Chayvialle, J.A., and Cuber, J.C. (1998). Effects of neurotransmitters, gut hormones, and inflammatory mediators on mucus discharge in rat colon. Am J Physiol 275, G1073–1084.

Pogorzala, L.A., Mishra, S.K., and Hoon, M.A. (2013). The cellular code for mammalian thermosensation. J Neurosci 33, 5533–5541.

Rao, M., Nelms, B.D., Dong, L., Salinas-Rios, V., Rutlin, M., Gershon, M.D., and Corfas, G. (2015). Enteric glia express proteolipid protein 1 and are a transcriptionally unique population of glia in the mammalian nervous system. Glia 63, 2040–2057.

Rao, M., Rastelli, D., Dong, L., Chiu, S., Setlik, W., Gershon, M.D., and Corfas, G. (2017). Enteric Glia Regulate Gastrointestinal Motility but Are Not Required for Maintenance of the Epithelium in Mice. Gastroenterology 153, 1068–1081 e1067.

Rescigno, M., Urbano, M., Valzasina, B., Francolini, M., Rotta, G., Bonasio, R., Granucci, F., Kraehenbuhl, J.P., and Ricciardi-Castagnoli, P. (2001). Dendritic cells express tight junction proteins and penetrate gut epithelial monolayers to sample bacteria. Nat Immunol 2, 361–367.

Rios, D., Wood, M.B., Li, J., Chassaing, B., Gewirtz, A.T., and Williams, I.R. (2016). Antigen sampling by intestinal M cells is the principal pathway initiating mucosal IgA production to commensal enteric bacteria. Mucosal Immunol 9, 907–916.

Rivera-Chavez, F., Lopez, C.A., Zhang, L.F., Garcia-Pastor, L., Chavez-Arroyo, A., Lokken, K.L., Tsolis, R.M., Winter, S.E., and Baumler, A.J. (2016). Energy Taxis toward Host-Derived Nitrate Supports a Salmonella Pathogenicity Island 1-Independent Mechanism of Invasion. MBio 7.

Sansonetti, P.J. (2004). War and peace at mucosal surfaces. Nat Rev Immunol 4, 953–964.

Sansonetti, P.J., and Phalipon, A. (1999). M cells as ports of entry for enteroinvasive pathogens: mechanisms of interaction, consequences for the disease process. Semin Immunol 11, 193–203.

Santos, R.L., Raffatellu, M., Bevins, C.L., Adams, L.G., Tukel, C., Tsolis, R.M., and Baumler, A.J. (2009). Life in the inflamed intestine, Salmonella style. Trends Microbiol 17, 498–506.

Sato, S., Kaneto, S., Shibata, N., Takahashi, Y., Okura, H., Yuki, Y., Kunisawa, J., and Kiyono, H. (2012). Transcription factor Spi-B–dependent and –independent pathways for the development of Peyer’s patch M cells. Mucosal Immunology 6, 838.

Sato, T., Vries, R.G., Snippert, H.J., van de Wetering, M., Barker, N., Stange, D.E., van Es, J.H., Abo, A., Kujala, P., Peters, P.J., et al. (2009). Single Lgr5 stem cells build crypt-villus structures in vitro without a mesenchymal niche. Nature 459, 262–265.

Savidge, T.C., Smith, M.W., James, P.S., and Aldred, P. (1991). Salmonella-induced M-cell formation in germ-free mouse Peyer’s patch tissue. Am J Pathol 139, 177–184.

Schierack, P., Rodiger, S., Kolenda, R., Hiemann, R., Berger, E., Grzymajlo, K., Swidsinski, A., Juretzek, T., Meissner, D., Mydlak, K., et al. (2015). Species-specific and pathotype-specific binding of bacteria to zymogen granule membrane glycoprotein 2 (GP2). Gut 64, 517–519.

Sharkey, K.A., Beck, P.L., and McKay, D.M. (2018). Neuroimmunophysiology of the gut: advances and emerging concepts focusing on the epithelium. Nat Rev Gastroenterol Hepatol 15, 765–784.

Sharkey, K.A., and Savidge, T.C. (2014). Role of enteric neurotransmission in host defense and protection of the gastrointestinal tract. Auton Neurosci 181, 94–106.

Snoek, S.A., Borensztajn, K.S., van den Wijngaard, R.M., and de Jonge, W.J. (2010). Neuropeptide receptors in intestinal disease: physiology and therapeutic potential. Curr Pharm Des 16, 1091–1105.

Stecher, B., and Hardt, W.D. (2011). Mechanisms controlling pathogen colonization of the gut. Curr Opin Microbiol 14, 82–91.

Stecher, B., Macpherson, A.J., Hapfelmeier, S., Kremer, M., Stallmach, T., and Hardt, W.D. (2005). Comparison of Salmonella enterica serovar Typhimurium colitis in germfree mice and mice pretreated with streptomycin. Infect Immun 73, 3228–3241.

Sudhof, T.C. (2012). Calcium control of neurotransmitter release. Cold Spring Harb Perspect Biol 4, a011353.

Szolcsanyi, J., Szallasi, A., Szallasi, Z., Joo, F., and Blumberg, P.M. (1990). Resiniferatoxin: an ultrapotent selective modulator of capsaicin-sensitive primary afferent neurons. Journal of Pharmacology and Experimental Therapeutics 255, 923–928.

Szolcsanyi, J., Szallasi, A., Szallasi, Z., Joo, F., and Blumberg, P.M. (1991). Resiniferatoxin. An ultrapotent neurotoxin of capsaicin-sensitive primary afferent neurons. Ann N Y Acad Sci 632, 473–475.

Tahoun, A., Mahajan, S., Paxton, E., Malterer, G., Donaldson, D.S., Wang, D., Tan, A., Gillespie, T.L., O’Shea, M., Roe, A.J., et al. (2012). Salmonella transforms follicle-associated epithelial cells into M cells to promote intestinal invasion. Cell Host Microbe 12, 645–656.

Takahashi, N., Matsuda, Y., Sato, K., de Jong, P.R., Bertin, S., Tabeta, K., and Yamazaki, K. (2016). Neuronal TRPV1 activation regulates alveolar bone resorption by suppressing osteoclastogenesis via CGRP. Sci Rep 6, 29294.

Takakura, I., Miyazawa, K., Kanaya, T., Itani, W., Watanabe, K., Ohwada, S., Watanabe, H., Hondo, T., Rose, M.T., Mori, T., et al. (2011). Orally administered prion protein is incorporated by m cells and spreads into lymphoid tissues with macrophages in prion protein knockout mice. Am J Pathol 179, 1301–1309.

Talbot, S., Abdulnour, R.E., Burkett, P.R., Lee, S., Cronin, S.J., Pascal, M.A., Laedermann, C., Foster, S.L., Tran, J.V., Lai, N., et al. (2015). Silencing Nociceptor Neurons Reduces Allergic Airway Inflammation. Neuron 87, 341–354.

Talbot, S., Foster, S.L., and Woolf, C.J. (2016). Neuroimmunity: Physiology and Pathology. Annu Rev Immunol 34, 421–447.

Thiennimitr, P., Winter, S.E., and Baumler, A.J. (2012). Salmonella, the host and its microbiota. Curr Opin Microbiol 15, 108–114.

Thompson, B.J., Washington, M.K., Kurre, U., Singh, M., Rula, E.Y., and Emeson, R.B. (2008). Protective roles of alpha-calcitonin and beta-calcitonin gene-related peptide in spontaneous and experimentally induced colitis. Dig Dis Sci 53, 229–241.

Trankner, D., Hahne, N., Sugino, K., Hoon, M.A., and Zuker, C. (2014). Population of sensory neurons essential for asthmatic hyperreactivity of inflamed airways. Proc Natl Acad Sci U S A 111, 11515–11520.

Tropini, C., Moss, E.L., Merrill, B.D., Ng, K.M., Higginbottom, S.K., Casavant, E.P., Gonzalez, C.G., Fremin, B., Bouley, D.M., Elias, J.E., et al. (2018). Transient Osmotic Perturbation Causes Long-Term Alteration to the Gut Microbiota. Cell 173, 1742–1754 e1717.

Tsai, P.Y., Zhang, B., He, W.Q., Zha, J.M., Odenwald, M.A., Singh, G., Tamura, A., Shen, L., Sailer, A., Yeruva, S., et al. (2017). IL-22 Upregulates Epithelial Claudin-2 to Drive Diarrhea and Enteric Pathogen Clearance. Cell Host Microbe 21, 671–681 e674.

Uesaka, T., Young, H.M., Pachnis, V., and Enomoto, H. (2016). Development of the intrinsic and extrinsic innervation of the gut. Dev Biol 417, 158–167.

Van Der Zanden, E.P., Boeckxstaens, G.E., and de Jonge, W.J. (2009). The vagus nerve as a modulator of intestinal inflammation. Neurogastroenterol Motil 21, 6–17.

Vazquez-Torres, A., Jones-Carson, J., Baumler, A.J., Falkow, S., Valdivia, R., Brown, W., Le, M., Berggren, R., Parks, W.T., and Fang, F.C. (1999). Extraintestinal dissemination of Salmonella by CD18-expressing phagocytes. Nature 401, 804–808.

Veiga-Fernandes, H., and Pachnis, V. (2017). Neuroimmune regulation during intestinal development and homeostasis. Nat Immunol 18, 116–122.

Vergnolle, N., and Cirillo, C. (2018). Neurons and Glia in the Enteric Nervous System and Epithelial Barrier Function. Physiology (Bethesda) 33, 269–280.

Voehringer, D., Liang, H.E., and Locksley, R.M. (2008). Homeostasis and effector function of lymphopenia-induced “memory-like” T cells in constitutively T cell-depleted mice. J Immunol 180, 4742–4753.

Vulchanova, L., Casey, M.A., Crabb, G.W., Kennedy, W.R., and Brown, D.R. (2007). Anatomical evidence for enteric neuroimmune interactions in Peyer’s patches. J Neuroimmunol 185, 64–74.

Walters, W.A., Xu, Z., and Knight, R. (2014). Meta-analyses of human gut microbes associated with obesity and IBD. FEBS Lett 588, 4223–4233.

Williams, I.R., and Owen, R.L. (2015). M Cells Specialized Antigen Sampling Cells in the Follicle-Associated Epithelium. Mucosal Immunology (Fourth 1, 211–229.

Woolf, C.J., and Ma, Q. (2007). Nociceptors--noxious stimulus detectors. Neuron 55, 353–364.

Xu, Z.-Z., Kim, Y.H., Bang, S., Zhang, Y., Berta, T., Wang, F., Oh, S.B., and Ji, R.-R. (2015). Inhibition of mechanical allodynia in neuropathic pain by TLR5-mediated A-fiber blockade. Nature Medicine 21, 1326.

Yang, N.J., and Chiu, I.M. (2017). Bacterial Signaling to the Nervous System through Toxins and Metabolites. Journal of Molecular Biology 429, 587–605.

Yoo, B.B., and Mazmanian, S.K. (2017). The Enteric Network: Interactions between the Immune and Nervous Systems of the Gut. Immunity 46, 910–926.

Zeisel, A., Hochgerner, H., Lonnerberg, P., Johnsson, A., Memic, F., van der Zwan, J., Haring, M., Braun, E., Borm, L.E., La Manno, G., et al. (2018). Molecular Architecture of the Mouse Nervous System. Cell 174, 999–1014 e1022.

