## Supplementary figures and images for "Gut-innervating nociceptor neurons protect against enteric infection by modulating the microbiota and Peyer’s patch microfold cells"

### Supplementary Figures S1-S7

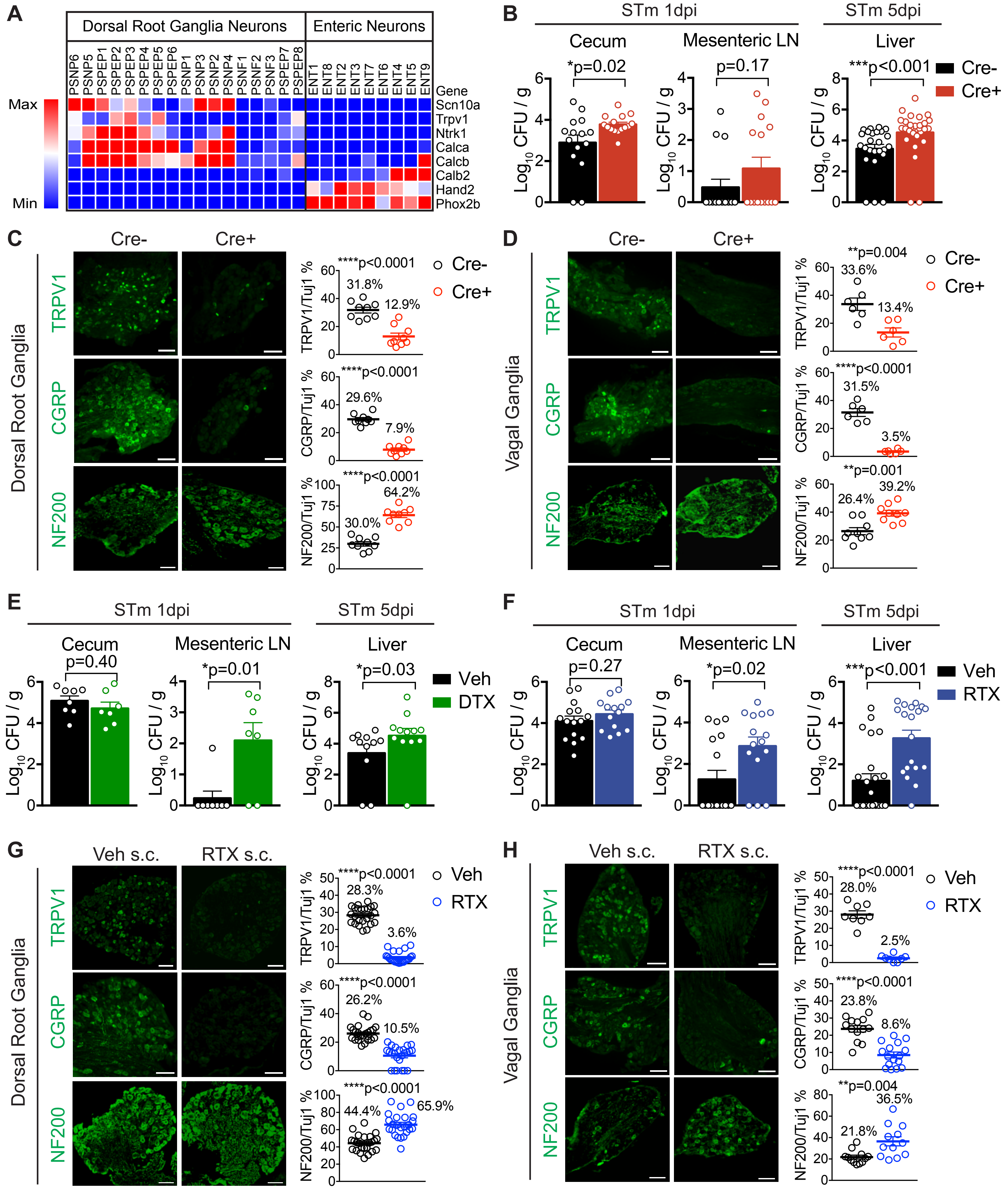

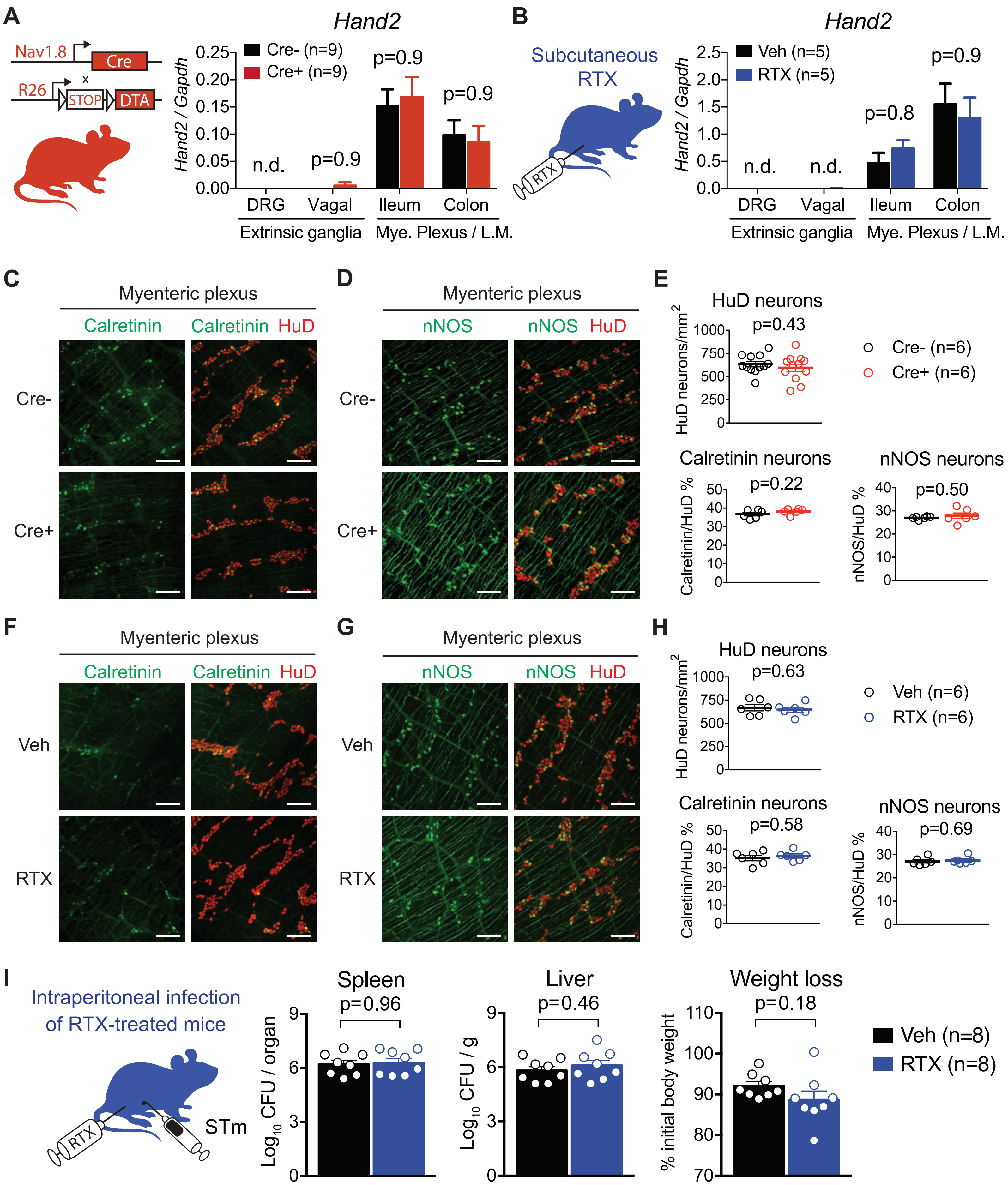

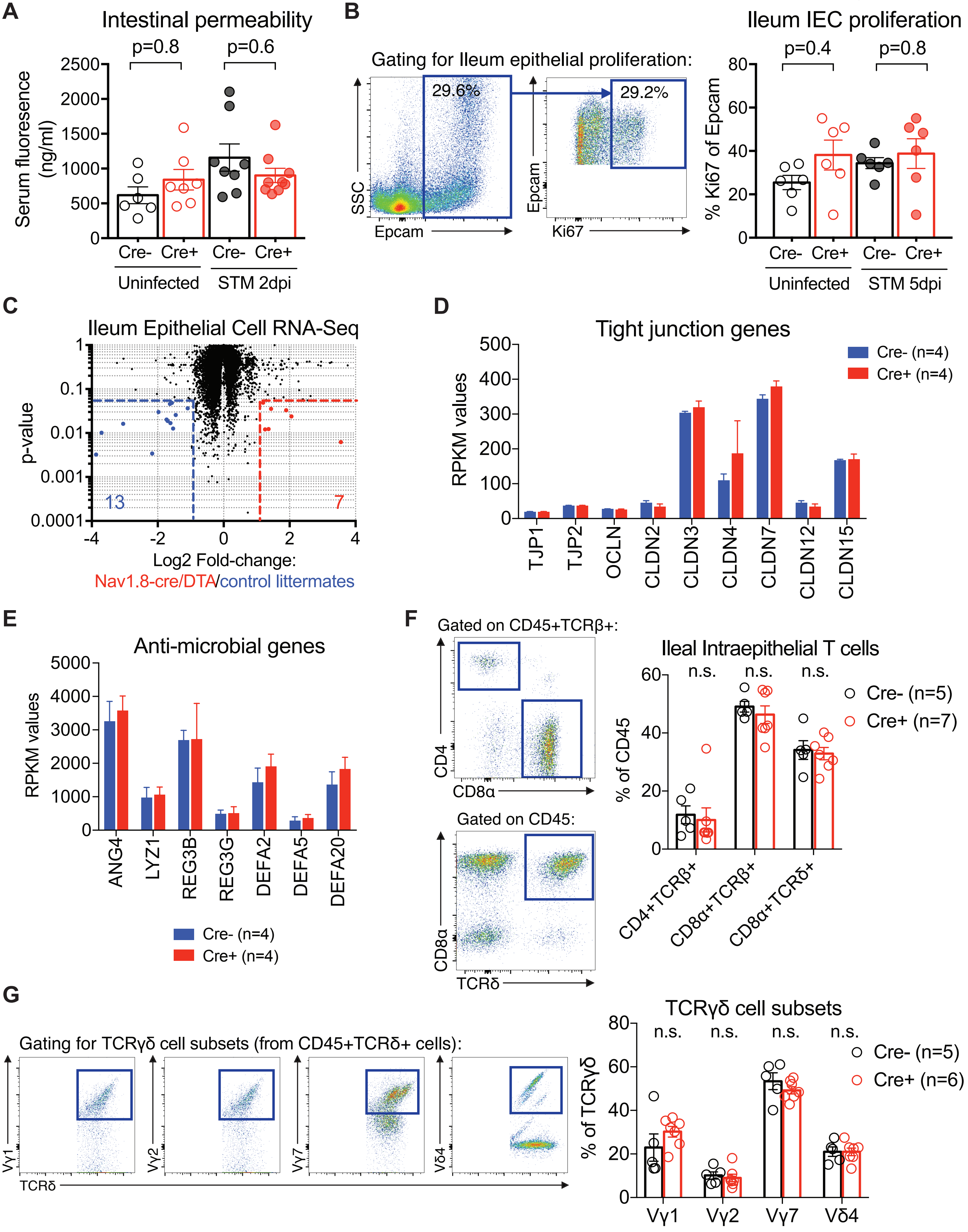

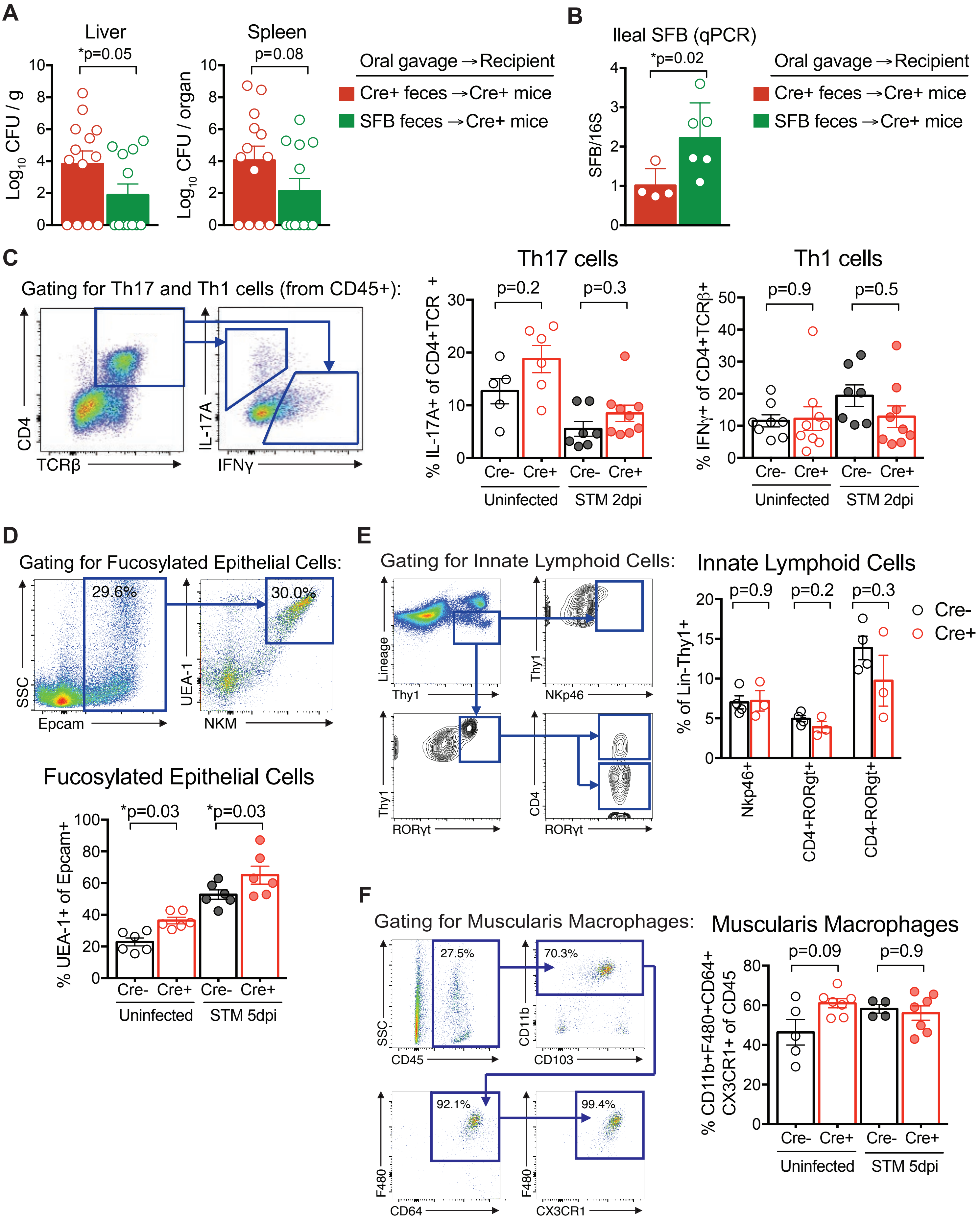

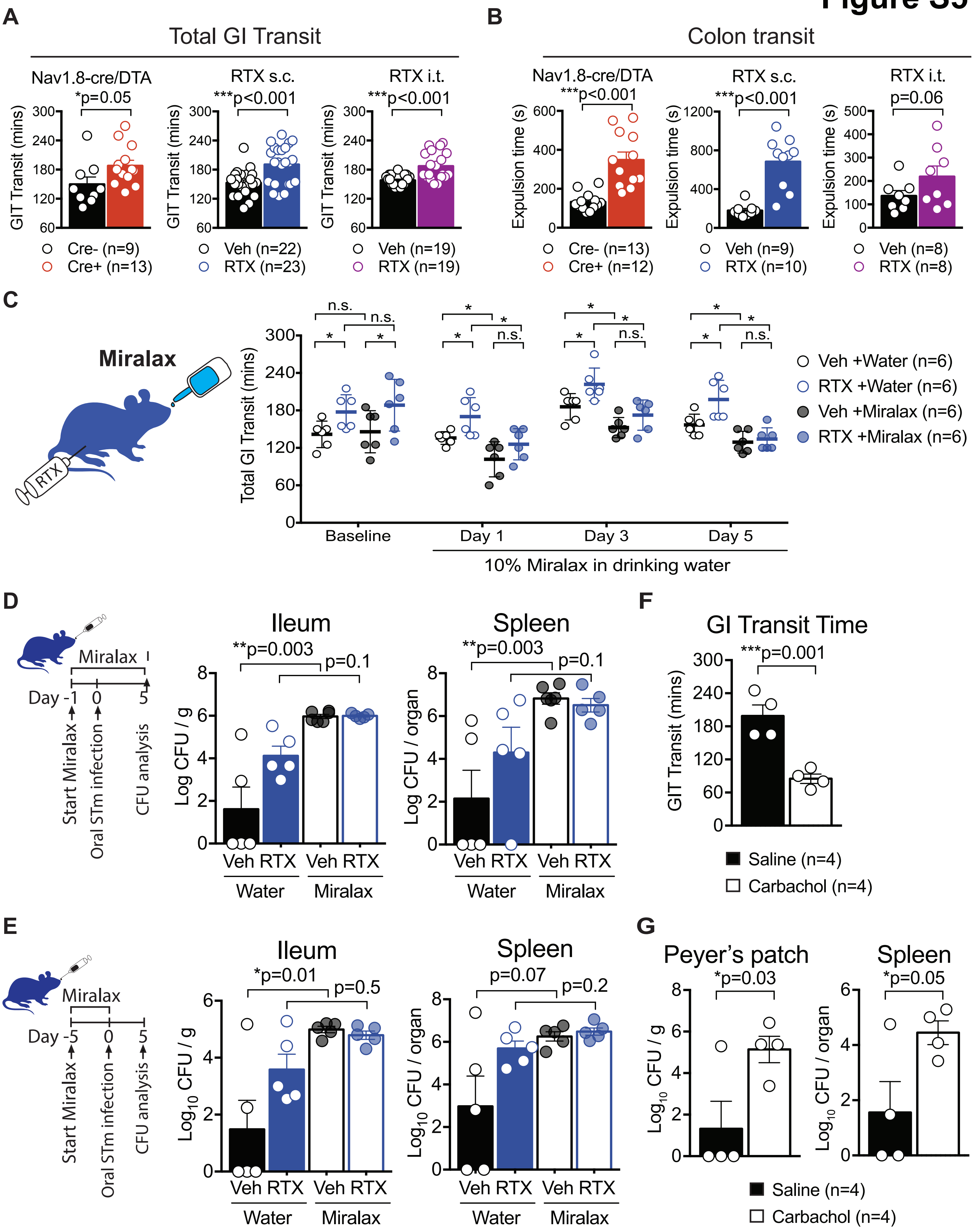

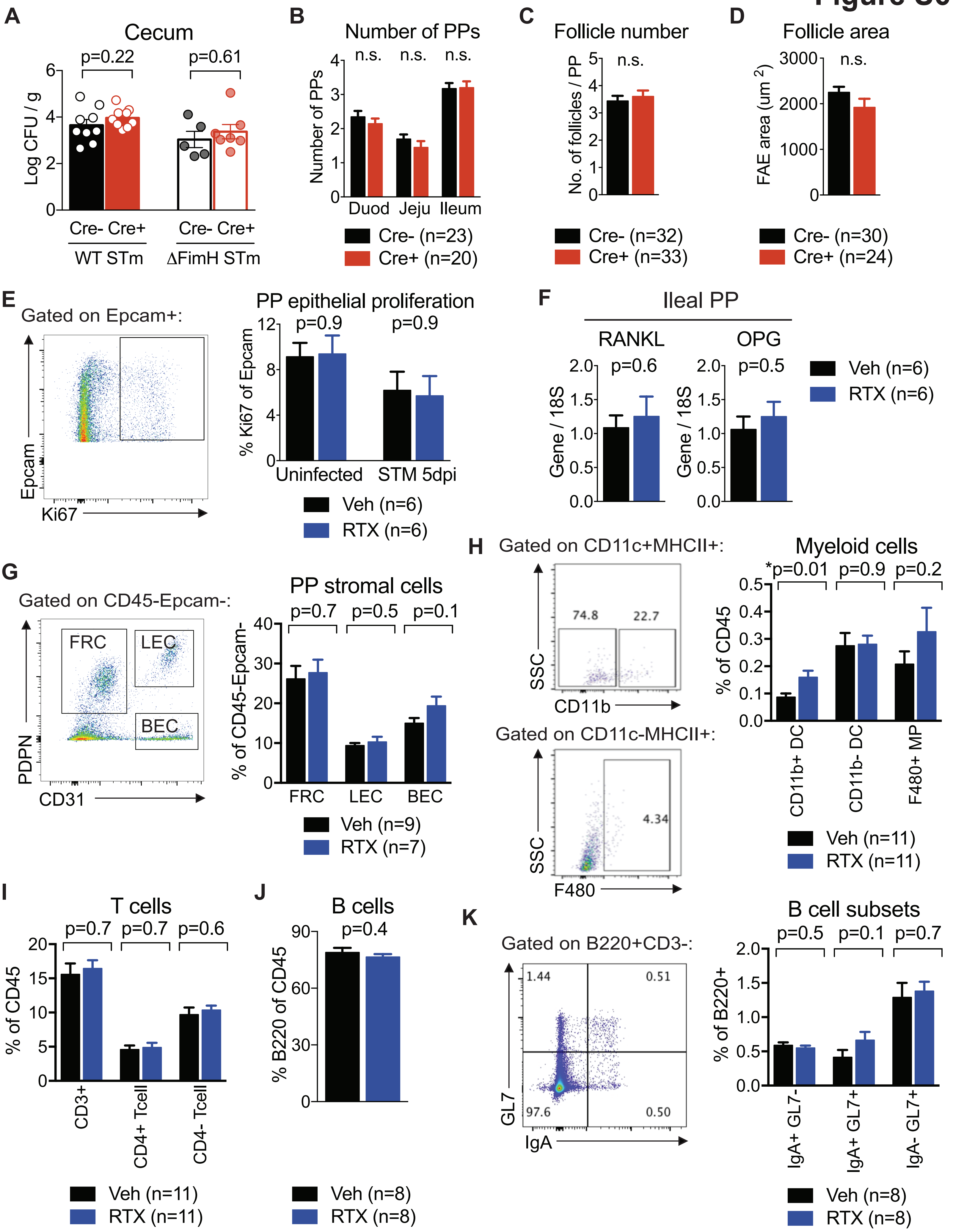

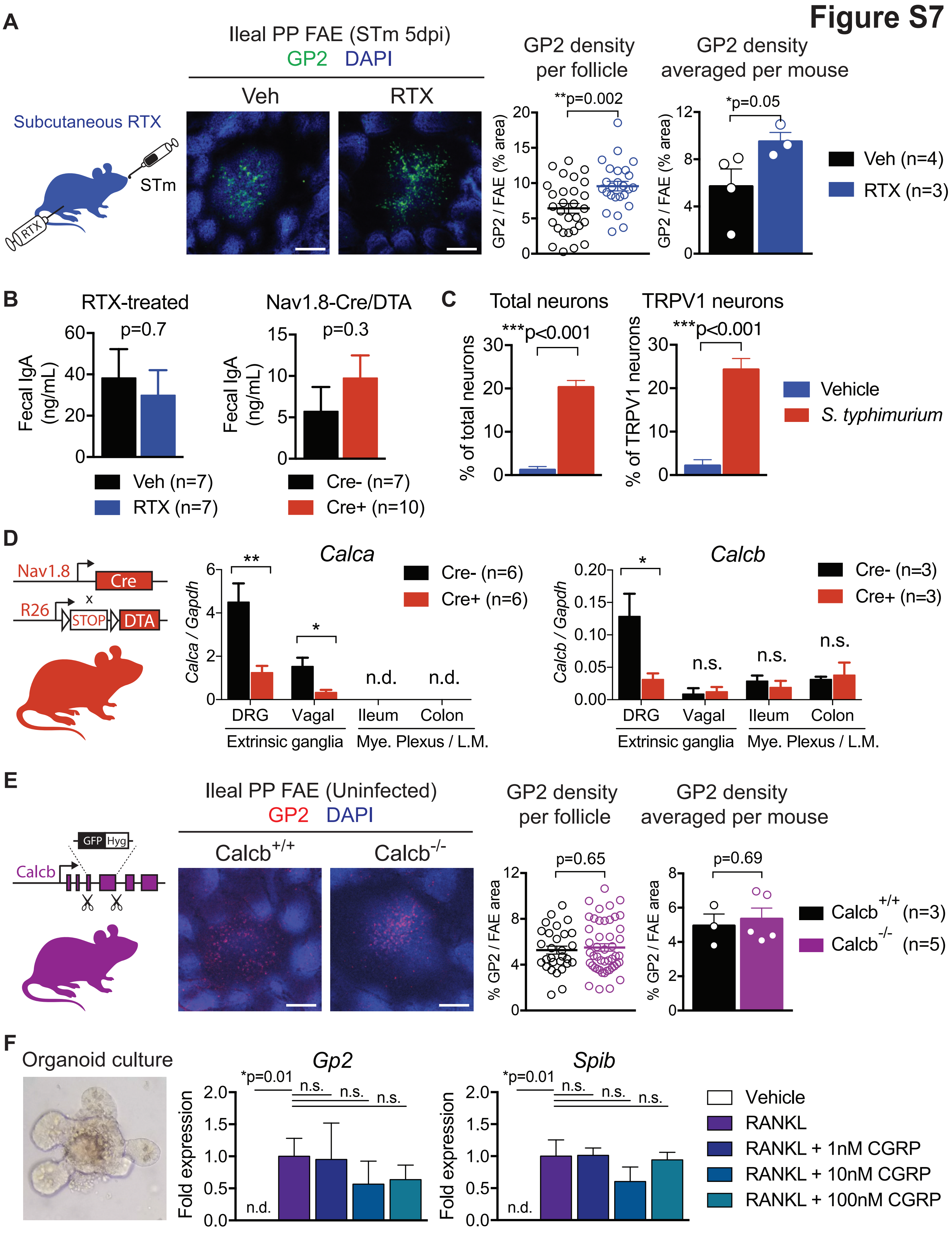
